# Comprehensive mutational characterization of the calcium-sensing STIM1 EF-hand reveals residues essential for structure and function

**DOI:** 10.1101/2025.01.23.634525

**Authors:** Nisha D. Kamath, Kenneth A. Matreyek

**Affiliations:** Department of Pathology, Case Western Reserve University School of Medicine, Cleveland, Ohio, USA

## Abstract

Calcium signaling is a fundamental molecular means of cellular regulation. Store operated calcium entry (SOCE) is a major intracellular signaling module, wherein calcium release from the endoplasmic reticulum triggers transmembrane STIM1 proteins to conformationally shift and oligomerize to prompt calcium influx from the extracellular environment. STIM1 senses ER calcium concentrations with its canonical EF-hand domain, and missense variants can dysregulate SOCE and cause Tubular Aggregate Myopathy, Stormorken Syndrome or immunodeficiency. Few STIM1 EF-hand variants are characterized, obscuring how STIM1 sequence controls its function, and hampering clinical interpretation of STIM1 variants observed in patients.

We leveraged fitness costs caused by overexpression of STIM1 variants in cultured human cells to functionally characterize 706 of the 720 possible single amino acid variants of the STIM1 canonical EF-hand. The calcium-coordinating EF-hand residues exhibited varying mutational patterns. The trailing helix possessed a core of immutable residues, even depleting during library propagation in bacteria, implicating residues normally restraining STIM1 aggregation. The leading helix only exhibited toxicity in cells with endogenous STIM1, implicating a multimerization-dependent STIM1 regulatory module. No cytotoxic STIM1 variants were observed in healthy human populations. Some disease-associated variants had low scores, but most pathogenic variants were not overtly cytotoxic in our assay. We demonstrate that orthogonal measurements for STIM1 oligomerization, cytoplasmic calcium influx, and cellular stress complement the cytotoxicity phenotypes to enhance variant understanding. Collectively, these data reveal the complex molecular roles embedded in the STIM1 canonical EF-hand sequence for its function in promoting calcium signaling through SOCE.

## Introduction

Engagement of various cell surface receptors – such as the B-cell receptors, T-cell receptors, and many G-protein coupled receptors – activate store operated calcium entry (SOCE), leading to a large influx of extracellular calcium ions into the cell cytoplasm. SOCE generally occurs when receptor activation triggers the release of secondary messenger molecules that cause an initial efflux of calcium out of the endoplasmic reticulum (ER) (Putney, 1986; Takemura et al., 1989). In non-excitable mammalian cells, this large decrease in ER calcium concentration triggers the activation of STIM proteins STIM1 (Roos et al., 2005) and STIM2 (Brandman et al., 2007), single pass ER transmembrane proteins that act as calcium “sensors” within the ER lumen (Liou et al., 2005) (**Figure 1A**). Once activated, STIM proteins oligomerize (Luik et al., 2008) and couple with ORAI family plasma membrane channel proteins (Prakriya et al., 2006), which open to allow calcium ions to flow down their concentration gradient into the cytosol (**Figure 1A**). There, the calcium ions bind various proteins including calmodulin and calcineurin, which modulate effector proteins to alter the cellular state in response to the extracellular stimuli, such as gene transcription. SOCE also plays roles in cellular homeostasis, such as refilling intracellular Ca^2+^ stores and supporting muscle contractility (Soboloff et al., 2012).

**Figure 1.**
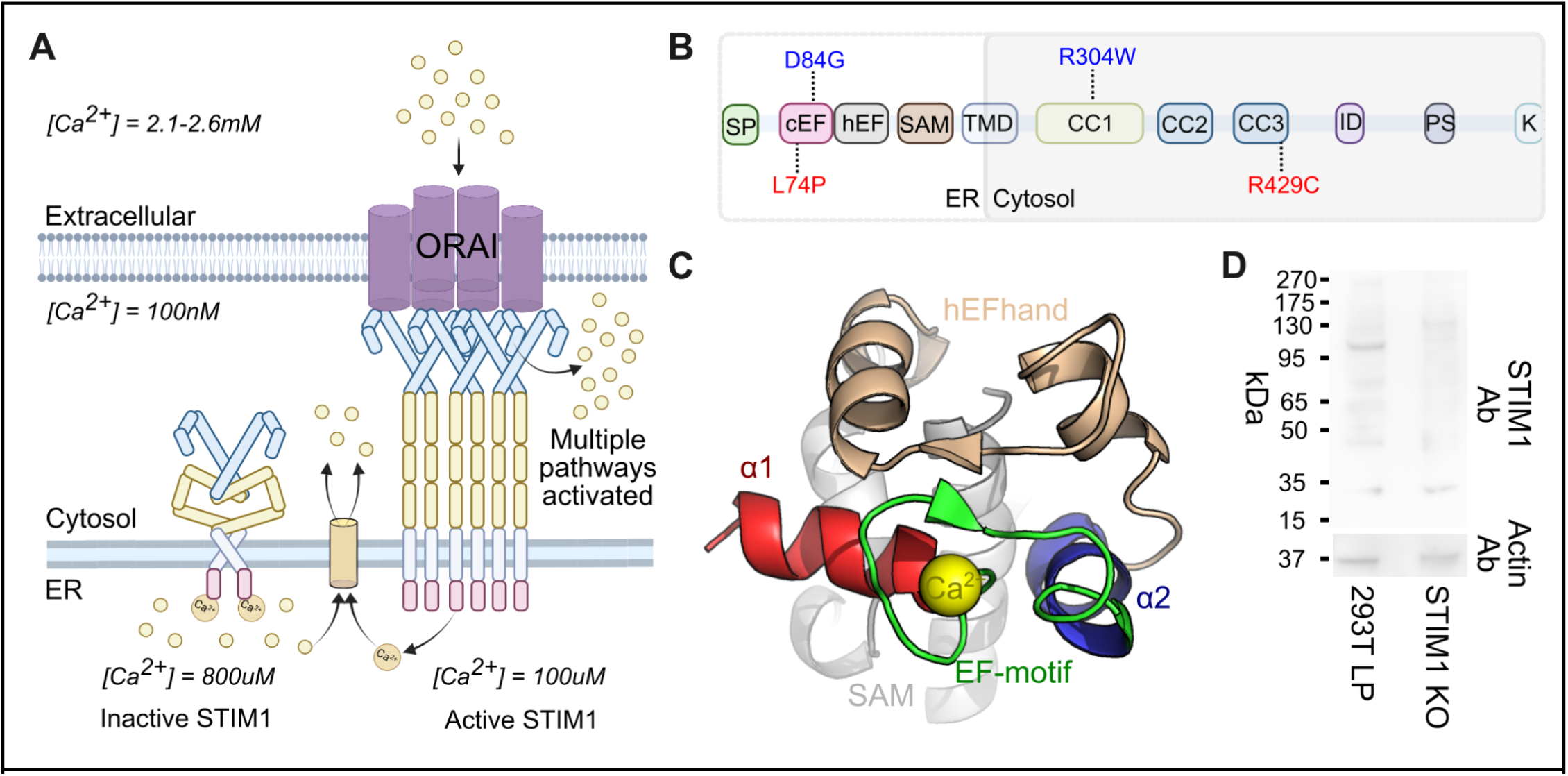
STIM1 calcium sensing with the cEF-hand during SOCE: A) Schematic representation of STIM1 during SOCE. B) Domains of STIM1 labeled with previously characterized variants used as controls for subsequent experiments. Blue are variants known to cause TAM or STRMK. Red are variants known to cause immunodeficiency. SP- signal peptide; cEF- canonical EF-hand; hEF - hidden EF-hand; SAM- sterile alpha motif; TMD- transmembrane domain; CC1- coiled-coil 1 domain; CC2- coiled-coil 2 domain; CC3- coiled-coil 3 domain; ID - inhibitory domain; PS- proline/serine-rich domain; K- lysine-rich domain. C) NMR structure of the STIM1 lumenal domain (PDB: 2K60 (Stathopulos et al., 2008)). D) Western blot using STIM1-targeting antibody to detect endogenously expressed STIM1 in standard and knockout HEK293T landing pad cells.

STIM proteins serve as ER calcium sensors by linking calcium binding at their ER luminal ends with conformational rearrangements at their cytoplasmic ends. STIM1 is the stronger activator of ORAI and is thought to play a dominant role in SOCE (Liou et al., 2005; Soboloff et al., 2012). When inactive, STIM1 forms a dimer through its cytosolic section. Calcium sensing is initiated by the ER-luminal section, which harbors a pair of EF-hand domains and a sterile alpha motif (SAM) domain. The STIM1 canonical EF-hand (cEF-hand) contains a central EF-hand motif, a 12 amino acid loop containing carboxyl and carbonyl oxygens required for coordination of a calcium ion (Clapham, 2007), bookended by alpha helices creating an overall helix-loop-helix (**Figure 1B,C**). The paired non-canonical “hidden” EF-hand (hEF-hand), which has evolutionarily lost the ability to bind calcium, supports proper folding of the lumenal domain (Huang et al., 2009; Stathopulos et al., 2008). This is unique to STIM molecules, as other calcium binding proteins like calmodulin typically bind calcium at all EF-hands present (Marshall et al., 2015).

The STIM1 cEF-hand has weak affinity for calcium (Stathopulos et al., 2006), and the release of calcium from the ER following receptor protein activation is sufficient to dissociate the bound calcium ion and trigger its conformational rearrangement, inducing aggregation of the ER-intraluminal EF-SAM domains, oligomerization of the STIM1 cytosolic extensions, and coupling with hexameric ORAI plasma membrane channels to permit large-scale calcium influx. Germline mutations in the STIM1 cEF-hand are associated with tubular aggregate myopathy (TAM) (Böhm et al., 2013), Stormorken syndrome (STRMK) (Stormorken et al., 1985), or immunodeficiency (Feske, 2010). Due to its unstable and aggregative nature, only the calcium-bound form of the isolated STIM1 EF-SAM has been resolved to atomic resolution (Stathopulos et al., 2008), and calcium-free conformations have been limited to general structural interpretations made with solution NMR (Enomoto et al., 2020; Sirko et al., 2022) and molecular dynamics models (Sallinger et al., 2020). Despite its physiological importance, only a handful of STIM1 EF-hand coding variants have been characterized, and the forms of molecular dysregulation occurring with each disease-associated variant are unknown. A comprehensive, unbiased analysis of each residue within the cEF-hand domain is needed but has not yet been undertaken.

In this study, we systematically assessed how overexpression of every possible STIM1 cEF-hand missense variant impacted the survival of cultured human cells. By comparing our findings with existing structural data, we identified key residues regulating STIM1 structure and function. We demonstrated that STIM1 variants that exhibit low survival in our assay correspond to variants associated with disease in people. This large-scale functional analysis of the STIM1 cEF-hand domain yielded new insights into the complex relationships between STIM1 sequence, structure, and function, and provided functional evidence aiding our understanding of how STIM1 dysregulation alters SOCE and can contribute to SOCE-related pathogenesis in humans.

## Results

### Developing a cell-survival assay for the functional characterization of STIM1 variants

STIM1 multimerization is a key step in its cellular function, and the cellular impacts of any given STIM1 variant may be influenced by the presence of an additional WT copy in the same cell. We thus performed our STIM1 studies both in the presence and absence of endogenous STIM1. Standard LP cells were HEK293T landing pad cells with endogenous STIM1 left intact, from which we generated knockout LP cells through biallelic deletion of exon 2 of the STIM1 gene located on chromosome 11. This created a frameshift at residue 47 in the resulting STIM1 open reading frame and the presumed absence of any possible full-length protein. Western blot analysis confirmed the successful knockout of STIM1 (**Figure 1D**).

To characterize the effect of nearly all possible single amino acid variants on STIM1 cEF-hand function, we harnessed a cell manipulation format compatible with pooled genetic assays for protein variant activities. We thus utilized engineered “landing pad” HEK 293T cells (LP) that encode a singular genomically integrated attP recombination site for the Bxb1 serine recombinase, preceded by a Tet inducible promoter (Matreyek et al., 2017). This clonally derived cell line can then be transfected with a library of transgenic plasmids encoding a complementary attB Bxb1 recombination site. DNA recombination between the attB and attP sites results in irreversible, directional insertion of a singular plasmid molecule into the chromosomally embedded landing pad site, leading to a single variant from the pooled variant library being expressed per cell (**Figure 2A**). This transgenic expression format provides a strict genotype-phenotype link needed for multiplex assays with pooled plasmid libraries encoding a wide range of protein-coding variants.

**Figure 2.**
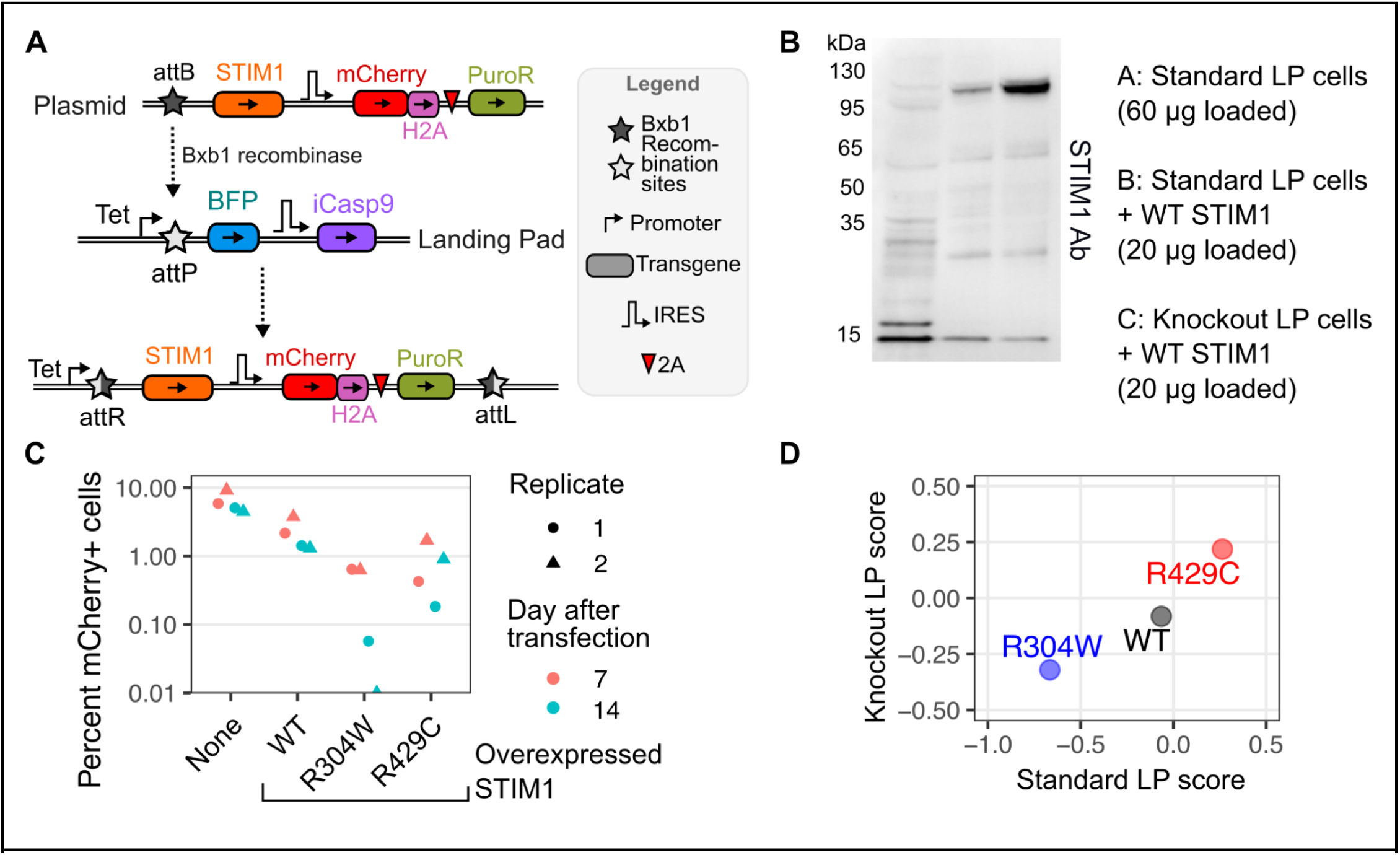
Impact of single-copy STIM1 cDNA overexpression on cell survival: A) Schematic of recombination of the plasmid construct into the landing pad locus. B) Western blot against STIM1 for standard LP cells, standard LP cells with STIM1 overexpression, and knockout LP cells with STIM1 overexpression. C) Depletion of STIM1 and mCherry expressing cells from day 7 to day 14 after transfection. D) Scatterplot of survival scores of pooled STIM1 control variants in both standard LP and knockout LP conditions.

Overexpressing the reference (WT) human STIM1 protein at high levels from the landing pad was toxic to the cells. Cells encoding the human STIM1 cDNA preceded by a consensus translation initiation sequence rapidly depleted from culture compared to cells engineered with a control construct (**figure supplement 1A**). We altered the translation initiation sequence to one that reduces steady-state protein levels roughly 50-fold (Shukla et al., 2021). Despite the cells no longer depleting from culture (**figure supplement 1A**), the STIM1 protein produced from the landing pad locus was substantially higher than endogenously present in these HEK 293T cells (**Figure 2B – figure supplement 1B-C**). Knowing that STIM1 dysregulation, such as its gross overexpression, could elicit a survival defect in cultured cells, we explored its potential as a phenotypic assay, and expressed all STIM1 variants under these expression conditions for the entirety of the study.

Mysregulated STIM1 activation causes abnormally high intracellular calcium influx that can overload the cell and initiate apoptosis (Tanwar and Motiani, 2018). The autosomally dominant, Stormorken syndrome variant STIM1 R304W causes high calcium influx through spontaneous activation of ORAI1 in the absence of ER calcium depletion (Misceo et al., 2014). In contrast, individuals homozygous for STIM1 R429C experience immunodeficiency and low calcium influx within the cell, as they lack STIM1 protein capable of coupling with ORAI channels (Fuchs et al., 2012). To assess our ability to phenotypically differentiate WT-like STIM1 variants from STIM1 variants causative for TAM/STRMK or immunodeficiency, we separately introduced WT STIM1, along with the R304W and R429C variants, into standard LP cells. Over the course of a week, we observed strong depletion of the R304W variant (**Figure 2C**), suggesting that these expression conditions were capable of distinguishing the TAM/STRMK causing disease variants from WT.

Pooled assay experiments can yield differing results from traditional arrayed experiments in unexpected ways, so we performed a preliminary pooled assay with the aforementioned R304W and R429C STIM1 variants. We sequenced the STIM1 variant library prior to and following integration into both the standard LP and knockout LP cells. Upon taking the frequency of DNA reads of each variant after integration, and dividing those values with the frequency of DNA reads of each variant in the plasmid mixture, we calculated a survival ratio. The survival ratio was log transformed to yield survival scores. Variants that depleted from the population yielded negative survival scores, whereas variants that survived yielded positive survival scores. There was clear phenotypic separation between variants in both standard and knockout conditions (**Figure 2D**). Based on these results, we proceeded with a deep mutational scan of the full 36-residue STIM1 cEF-hand domain to comprehensively assess the sequence-function relationships governing STIM1 calcium sensing.

### Deep mutational scanning of the STIM1 cEF-hand domain

We created a site-saturation mutagenesis library encoding every possible single amino acid variant of the STIM1 cEF-hand. The variant sequences were introduced into the human STIM1 cDNA with a tiled set of PCR primers, with each primer pair encoding a degenerate NNK codon for each of the 36 residues of the STIM1 cEF-hand (Jain and Varadarajan, 2014). The resulting library of plasmid molecules encoded 792 total missense, nonsense and synonymous variants of the STIM1 cEF-hand (**figure supplement 2A-C**).

We first recombined the plasmid library into standard LP cells. The STIM1 cEF-hand domain was amplified from the genomic DNA of recombined cells, sequenced, and scored 5 separate times (**Figure 3A – figure supplement 3A**). We obtained 744 variant-specific survival scores in standard LP cells (**Figure 3B**). Synonymous and nonsense variants typically serve as helpful controls in deep mutational scans, as synonymous variants typically exhibit WT-like effects, whereas nonsense variants do not. The synonymous variants exhibited a roughly normal distribution of scores (**Figure 3B, left**), and had the highest survival scores as a mutational class. In contrast, the distribution of scores for both missense and nonsense variants were skewed toward lower scores, suggesting that these classes included variants with survival defects. The mean score for the nonsense variants was comparable to the missense variants for this assay (**Figure 3B, left**).

**Figure 3.**
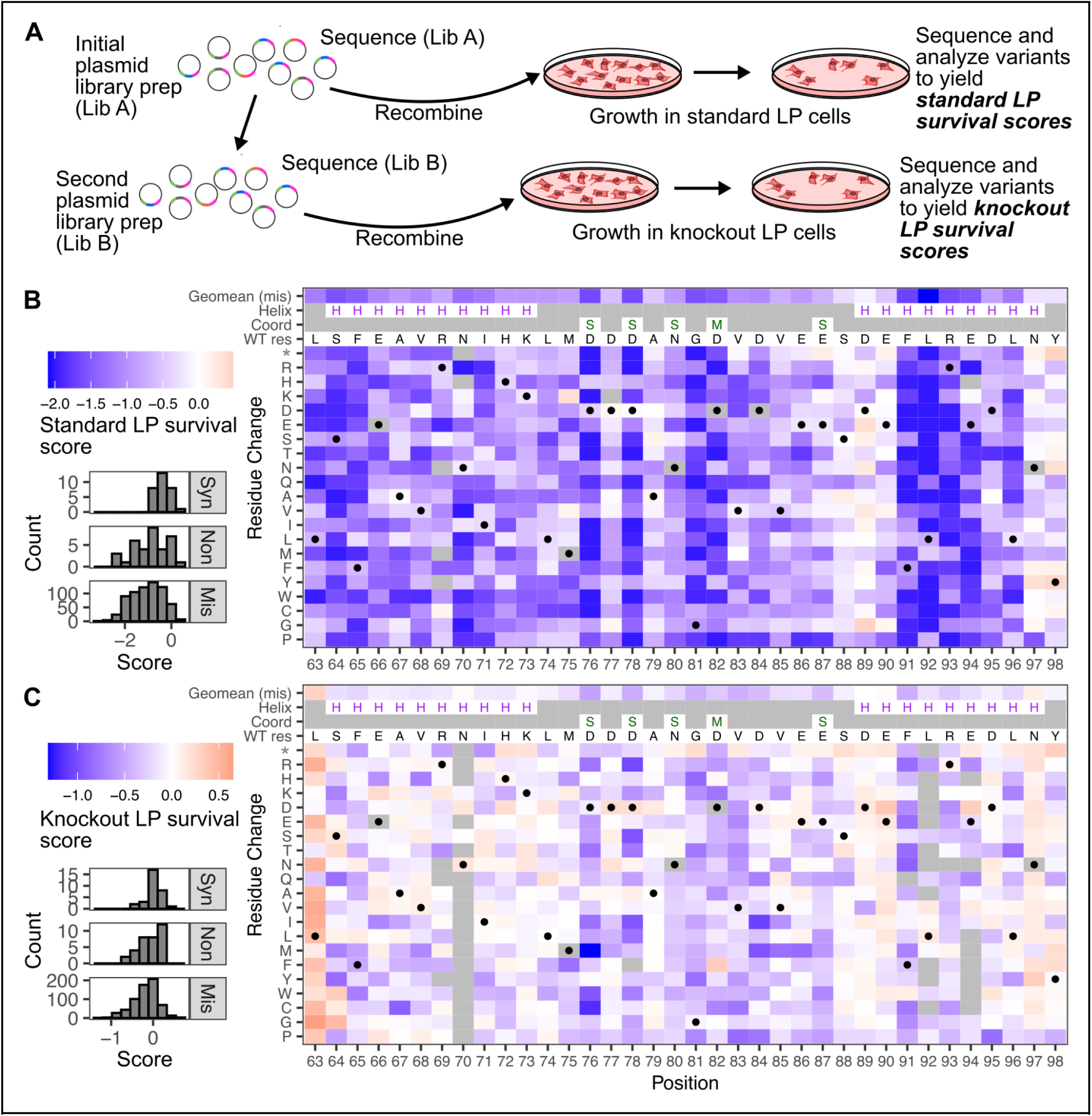
Cell surivival score maps of STIM1 cEF-hand variant overexpression: A) Schematic of the deep mutational scanning workflow. The variants in the plasmid library were recombined into standard LP cells. Cells were harvested 9 days following transfection. This process was repeated in knockout LP cells, but only after the original library (Lib A) was transformed into fresh bacteria to create more physical plasmid molecules (Lib B). B) Left: Histograms of scores for synonymous, nonsense, and missense variants, for standard LP cells. Right: Genotype-phenotype map of each single amino acid change in the STIM1 cEF-hand domain. Gray spaces indicate missing values. For ease of data representation, the heatmap color minimum was scaled to −2, with values smaller than −2 given the same dark blue color. WT STIM1 had a score of −0.06. H symbols represent residues with alpha-helical secondary structure in PDB 2K60 (Stathopulos et al., 2008). S and M symbols indicate side-chain and main-chain contacts with the calcium ion, respectively. C) The same plots as described in panel B, except for variants that were tested in knockout LP cells. WT STIM1 had a score of −0.05.

We subsequently recombined the plasmid library into knockout LP cells. The original plasmid library was first transformed into bacteria and re-extracted to yield sufficient plasmid material to enable this second round of cell culture experiments (**figure supplement 2A-C**). Upon recombination, the STIM1 cEF-hand domain was processed and analyzed identically as the experiments in standard LP cells, and repeated for a total of 6 independent replicates (**Figure 3A – figure supplement 3B**). We obtained 710 survival scores in the knockout LP cells, with 706 variants possessing scores in both cell backgrounds (**Figure 3C**). Similar to the original dataset collected in the standard LP cells, the synonymous variants exhibited the highest scores, while the missense and nonsense variants both skewed toward lower scores (**Figure 3C, left**). Despite following identical procedures, the magnitude of scores was exaggerated in standard LP cells as compared to the knockout LP cells (**figure supplement 3C**). This was consistent with the relative values observed in the pilot assay (**Figure 2D**).

To validate our survival scores, we performed a competition assay with a small set of selected variants (**figure supplement 3D-E**). For most variants, their scores relative to the rest of the set were similar between the two assays, in both cell backgrounds. L74P and E90* were exceptions, as they scored worse in the validation assays compared to the full deep mutational scans, regardless of cell background. D76* was also an outlier, although only within knockout LP cells. These results suggest that there may be context-specific factors that impact the cell depletion effect, at least for a subset of STIM1 cEF-hand variants.

### Unexpected positional depletion of variants in bacterial cells

We noticed that there were residues with variants scored in the standard LP dataset but absent in the knockout LP dataset. Upon comparing the frequencies of the variants in the two library plasmid preps used to perform the DMS experiments in the different cell contexts, we observed that sizable fluctuations to variant frequencies occurred upon re-transformation and propagation of the library in bacteria (**Figure 4A**). Synonymous variants were the least impacted mutational class (**Figure 4B**), but were still slightly skewed toward lower scores. There was a strong underlying positional effect, as the nonsense variant scores highly correlated with the geometric mean of missense variant scores at each position (**Figure 4C**). The variants missing in the knockout LP cell experiments corresponded to the positions most depleted upon transformation, such as Asn70, Phe91, Leu92, Arg93, and Glu94 (**Figure 3C**, **Figure 4A**).

**Figure 4.**
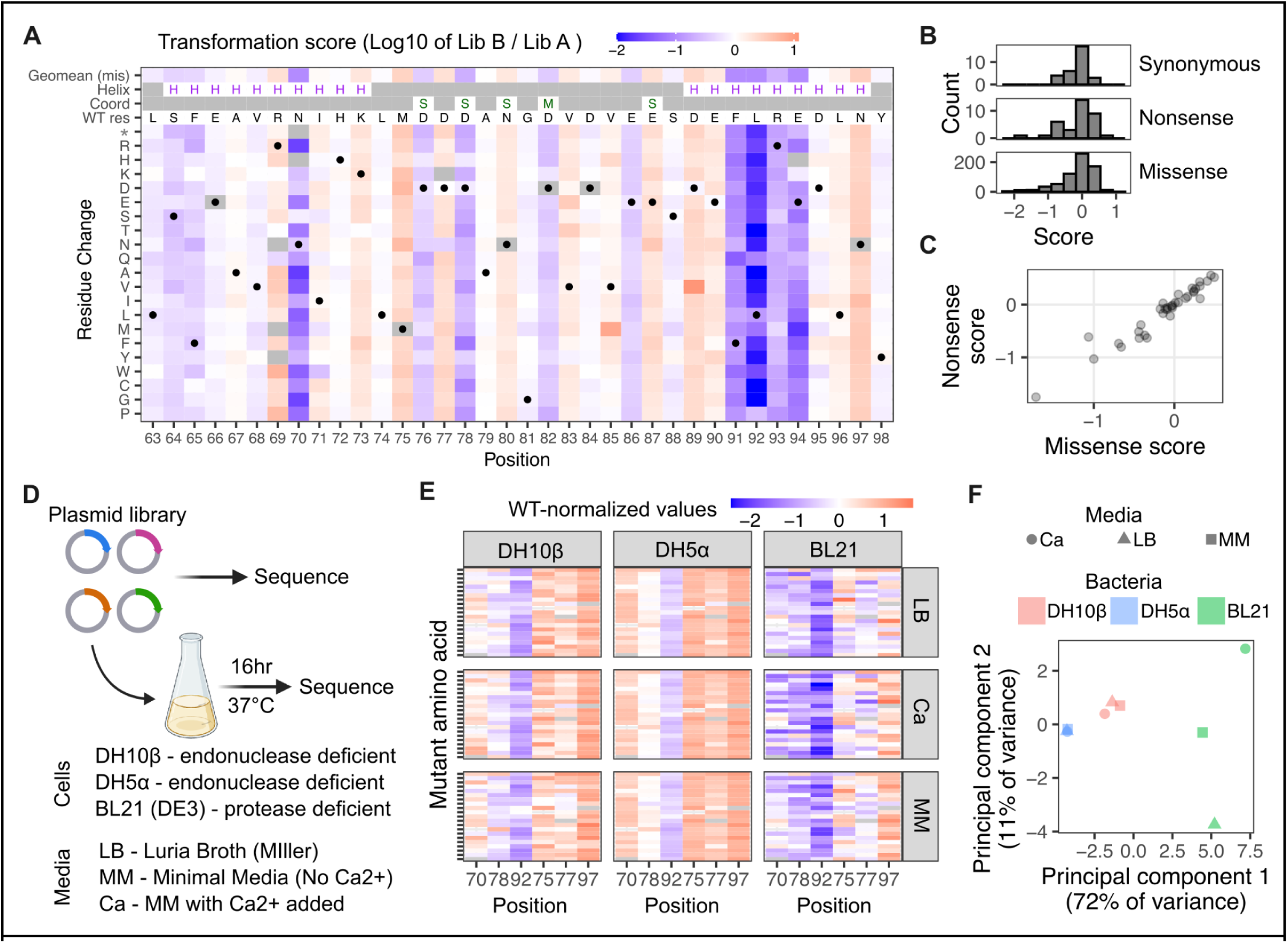
STIM1 cEF-hand variant depletion in bacteria: A) Genotype-phenotype map of variant-specific depletions from the plasmid library upon transformation. B) Histogram of missense, nonsense and synonymous variants for transformation scores. C) Scatterplot of nonsense variant and missense variant transformation scores averaged by position. D) Workflow schematic of experiments modulating bacterial strains and media. E) Genotype-phenotype maps and F) Principal component analysis of transformation scores grouped by bacterial strain and growth.

While originally unexpected, an investigation revealed an unannotated T7 promoter and nearby sigma factor pseudo binding sites, which may have driven low-level bacterial expression of the STIM1 coding sequence from our mammalian expression construct (**figure supplement 4A**). Western blotting of *E.coli* transformed with our mammalian STIM1 expression plasmid did not reveal expressed STIM1 protein (**figure supplement 4B**), although this approach is incapable of detecting partially translated STIM1 products.

To better understand the potential reasons underlying the observed effect, we performed a series of experiments modulating bacterial genetics and metabolic environment. We created a new mini-library of three depleted positions and three enriched positions, which were mixed together in equimolar amounts and transformed into three different strains of bacteria (**Figure 4D**). *E.coli* DH10β and DH5α are endonuclease deficient and widely used for plasmid transformation, while BL21(DE3) are protease deficient and typically used for heterologous protein expression. Furthermore, these particular BL21 cells express T7 polymerase, which will initiate transcription from the T7 promoter and further enhance any transcriptional and translational effects. Each transformed bacterial strain was grown in three different kinds of media: Luria Broth (LB), M9 minimal media (MM), and M9 minimal media with calcium added (Ca). This mini-library was independently transformed into the different *E.coli* strains and conditions three separate times, and each compared to the original plasmid library counts to calculate repeat transformation scores (**Figure 4D**).

Transformation into DH10β *E.coli*, which was the strain used to perform all previous transformations, reproduced the majority of the effects. There was strong correlation between the variant-specific scores in the full deep mutational scan as compared to this small-scale transformation (R^2^ of 0.72; **figure supplement 4C**). Transformation into DH5α *E.coli* looked similar to DH10β, except for a more pronounced positional bias (**Figure 4E**). Transformation into BL21 *E.coli* yielded the greatest magnitude of differences in variant-specific phenotypic effect. BL21 was also the only genetic background where the growth media substantially impacted phenotypic outcome (**Figure 4F**). Thus, at least part of the observed plasmid depletions were likely caused by fitness disadvantages experienced by bacteria expressing and translating STIM1 protein fragments.

### Comparison of variant effects across cell conditions

With three datasets of STIM1 cEF-hand sequence-function in hand, we assessed how each dataset compared to each other (**Figure 5A**). Both the standard LP and transformation scores had strong positional biases, and these positional effects correlated between the datasets (**Figure 5A**). Clustering of positionally averaged scores showed distinct subsets of STIM1 residues (**Figure 5B**). This analysis also identified residues that yielded low-scoring missense variants across cell conditions. These eight residues included a set of four contiguous amino acids, Phe91, Leu92, Arg93, and Glu94 (FLRE), with variants of Leu92 particularly toxic (**Figure 5B**).

**Figure 5.**
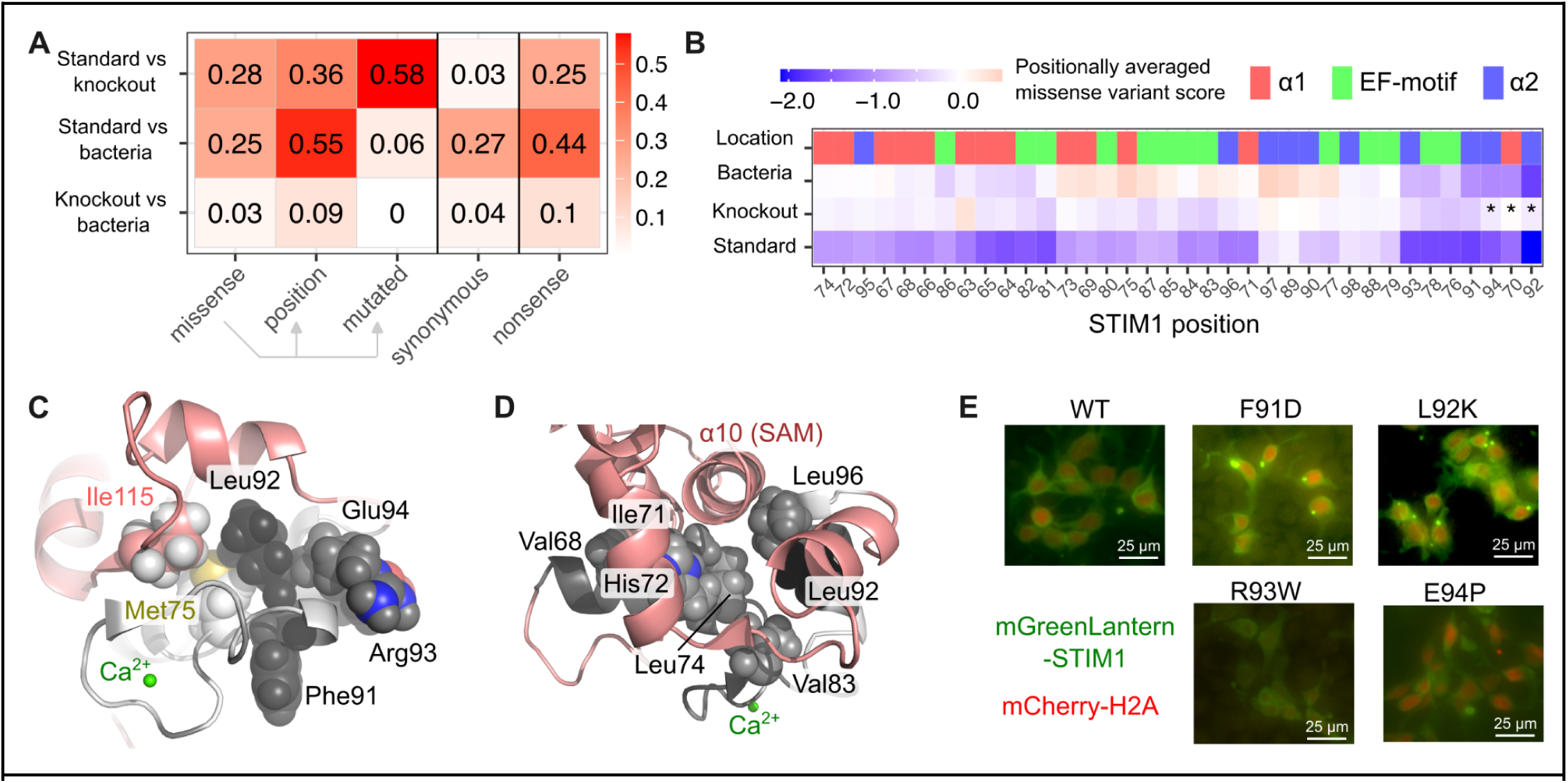
The central residues of helix α2 are critical for STIM1 function: A) Table of Spearman’s ρ^2^ values calculated upon comparison between data subsets across cell conditions. B) Heatmap of hierarchically clustered positionally averaged scores across cell conditions. Asterisks indicate positional averages heavily skewed by variant loss in bacteria. C) Pymol structure of cEF-hand showing the position of the ‘FLRE’ stretch of residues within helix α2. D) Pymol structure of cEF-hand residues involved in stabilizing the ER lumenal domains of STIM1 through binding to helix α10 located in the SAM domain. E) Fluorescent images of mGreenLantern-tagged STIM1 variants. The cell nucleus is demarcated by mCherry-H2A fluorescence.

This stretch of residues corresponds to the core of STIM1 helix α2 which is believed to unfold upon calcium release (Enomoto et al., 2020). Molecular dynamics simulations suggest that Leu92, which forms hydrophobic contacts with the hEF-hand and SAM domains in the calcium-bound state, experiences the greatest movement of all residues in the STIM1 EF-SAM domain once calcium is released (Sallinger et al., 2020; Schober et al., 2019). EF-hand domains exhibit a mixture of hydrophobic and charged amino acids at this C-terminal helix, which typically forms the more dynamic side of the EF-hand hydrophobic core (Denessiouk et al., 2014). In the available calcium-bound STIM1 cEF-hand NMR structure, Leu92 makes hydrophobic contacts with Met75 in the calcium-coordinating loop, along with Ile115 extending from the beta-strand on the hEF-hand (**Figure 5C**). Altogether, the toxicity introduced by coding substitutions at these residues support their importance in maintaining the proper structure of the STIM1 cEF-hand. Contrasingly, variants of residues at the ends of helix α2, corresponding to Asp89, Glu90, Asn97, and Tyr98, exhibited the highest variant scores within the cEF-hand (**Figure 5B**).

Leu92 is also one of a half-dozen hydrophobic residues in the EF-hand (Val68, Ile71, His72, Leu74, Val83, and Leu96) known to form a hydrophobic cleft that binds the SAM domain in the presence of calcium, thus stabilizing the ER lumenal domain of STIM1 (Stathopulos et al., 2008) (**Figure 5D**). Each of these residues were partially depleted in the standard LP cells, but no other positions exhibited the magnitude of depletion as Leu92 (**Figure 5B**). Thus, the impact of variants of Leu92 on cell survival within our expression systems highlight its particular importance in cEF-hand domain folding and conformational rearrangement.

Due to the uniquely cytotoxic effects exhibited by variants of the FLRE motif, we suspected that mutation of these residues may cause drastic misfolding and aggregation of the resulting protein. To assess the aggregation phenotypes of these variants within their cell contexts, we microscopically examined cells expressing STIM1 N-terminally tagged with mGreenLantern, without the addition of any SOCE inducer. In contrast to the largely diffuse ER- and cell surface fluorescence exhibited by WT STIM1, many of the cells that survived expression of F91D and L92K variants exhibited perinuclear puncta indicative of aggregated protein. On the other hand, R93W and E94P variants had similar diffuse fluorescence as WT STIM1 (**Figure 5E - figure supplement 5A**).

### Analysis of the EF-hand calcium coordinating residues

Next we focused on the EF-hand motif, which is a set of twelve residues in the central loop of the cEF-hand domain that directly coordinate calcium (**Figure 6A**). A subset of residues within this motif exhibited variant-specific toxicity in both standard and knockout conditions (**Figure 3B**, **Figure 3C**). Asp76 and Asp78 are the first two aspartic acids within the canonical DxDxDG EF-hand motif (Huang et al., 2009), and the scores of variants at these positions highly correlated between datasets (**Figure 6B**). Asn80 is a non-consensus amino acid for this motif (DxDxNG), and it was the calcium-coordinating residue most tolerant to substitution within STIM1 (**Figure 4B**). Expectedly, the N80D reversion to the canonical EF-motif amino acid had a high score. Glu87, which is the calcium coordinating residue furthest from the others, exhibited highly correlated albeit relatively small effects in both conditions.

**Figure 6.**
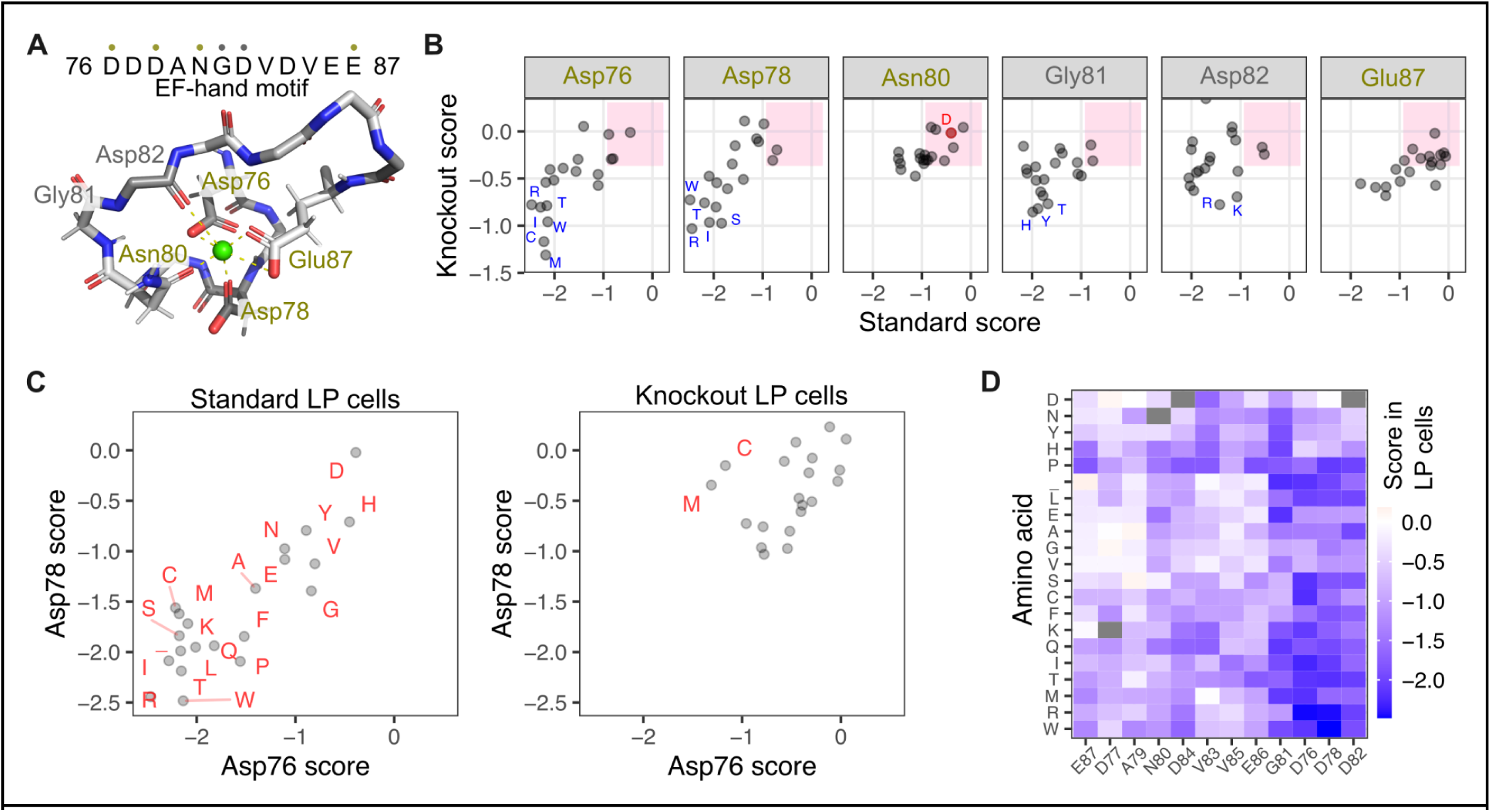
Analysis of the EF-hand calcium coordinating residues: A) NMR structure of the STIM1 cEF-hand motif (PDB: 2k60) with the peptide backbone and calcium coordinating side chain atoms shown as sticks. Hydrogen atoms are shown as thin sticks. The calcium ion is shown as a green sphere. B) Scatterplots comparing variant-specific scores across standard and knockout conditions for residues labeled in panel A. The pink box represents the 95 percent interval of synonymous variant scores,considered WT-like in survival. C) Scatterplot comparing mutation specific scores for Asp76 (x-axis) and Asp78 (y-axis), in standard LP cells (left) or knockout LP cells (right). D) Heatmap of EF-motif scores in standard LP cells, with positions ordered by hierarchical clustering based on mutation-specific tolerability, and mutations ordered by hierarchical clustering for similarities across positions.

Gly81 and Asp82 are involved in calcium coordination without their side chain atoms. Gly81 is critical for the tight turn needed by the carbon backbone of the EF-hand loop (**Figure 6A**). The rotational flexibility of Gly81 is also important for the proceeding carbonyl oxygen of Asp82 to coordinate calcium (Huang et al., 2009). Almost all variants of Gly81 and Asp82 exhibited low scores in standard LP cells, yet roughly half of these variants exhibited WT-like scores in knockout LP cells (**Figure 6B**). Thus, examination of the residues known to play direct roles in ion coordination revealed at least three groups differing in their magnitude effects across standard and knockout conditions.

We examined the EF-motif residues further by assessing their mutation-specific tolerability. Variants of Asp76 and Asp78 were highly correlated in their scores in standard LP cells (**Figure 6C, left**; Pearson’s R^2^: 0.75, Spearman’s ρ^2^: 0.54). The largest exceptions were cysteine and methionine, the two sulfur-containing amino acids. Asp76 and Asp78 exhibited reduced but observable correlation in knockout LP cells (**Figure 6C, right**). The difference in tolerability to cysteine and methionine were more pronounced in knockout conditions, with D76C and D76M far more detrimental to cell fitness than D78C or D78M. Clustering of EF-motif variants by residue demonstrated that Asp82 exhibited a similar mutation-specific tolerability as Asp78, and by extension, Asp76, in standard LP cells (**Figure 6D**). This correlation was lost in knockout conditions. Despite aspartate and glutamate side chains encoding seven of the twelve residues of the STIM1 cEF-motif, our mutational results reveal distinct functional roles played by distinct subsets of these residues depending on their location in the EF-motif.

### Distinct mutability of the cEF-hand entering helix

As a group, variants of residues in helix α1 exhibited relatively higher scores in knockout LP cells as compared to standard LP cells (**Figure 7A**, **Figure 7B**). Variants of Leu63, which directly precedes the structurally resolved helix α1, exhibited abnormally high scores in knockout LP cells (**Figure 7C**). In contrast, the EF-motif possessed a subset of variants that had lower scores in knockout LP cells as compared to the standard LP cells. These effects were attributable to variants of Val83, Asp84, and Glu87, which all lie on the latter-half of the EF-hand motif (**Figure 7C**).

**Figure 7.**
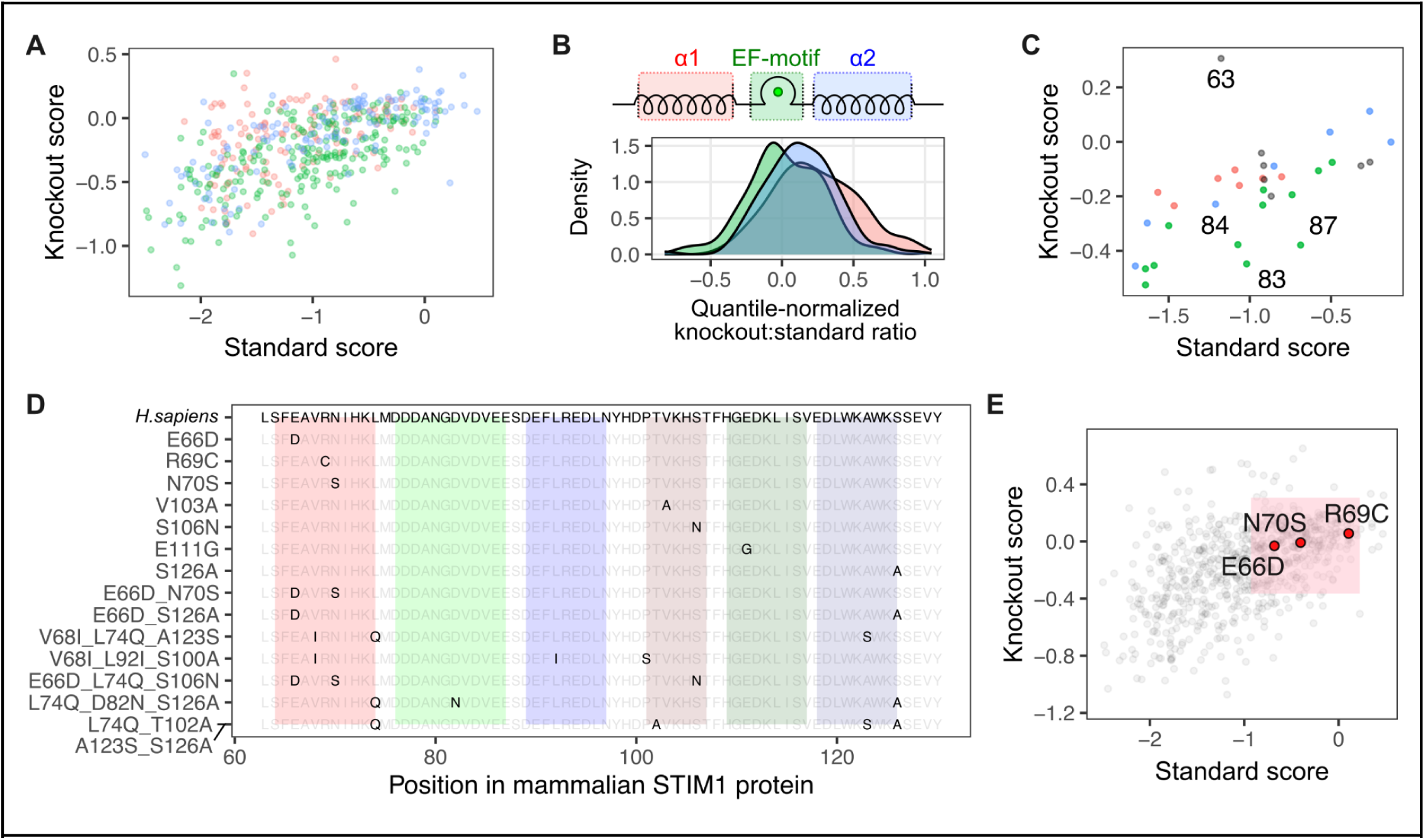
Sequence variability and context dependence of helix α1: A) Scatterplot of variant scores colored by presence in either helix α1 (red), the cEF-hand motif loop (green), or helix α2 (blue). B) Knockout to standard score ratio for variants grouped by section of the cEF-hand domain. C) Scatterplot of variant scores averaged on position and colored as in panel A. D) Sequence comparison of STIM1 cEF-hand domains found in mammalian orthologs E) Scatterplot of variant scores with scores of substitutions found in orthologs of STIM1 in red. The pink box represents the 95 percent interval of the synonymous variant scores, considered WT-like in survival.

Natural extant sequence variants highlight a distinct functional role for helix α1. Mammals exhibit minimal variation in the EF-motif and helix α2, but have substantially more variation in helix α1. The most common variants within mammals are Asp66 and Ser70, which are observed individually in other mammals, but also in combination, such as Asp66_Ser70 in felines (**Figure 7D**). R69C is present in the common marmoset. As expected, the corresponding E66D, R69C, and N70S variants of human STIM1 all had high scores in our assay (**Figure 7E**). This was particularly surprising for N70S, as the majority of other variants at this position had low scores. All three of these residues possess side chains that face outward from the same surface of helix α1 (**figure supplement 6A**). While indirect, these observations suggest that further investigation into a possible regulatory role for helix α1 in STIM1 oligomerization may be warranted.

### Interpretation of STIM1 EF-hand variants involved in human disease

We finally sought to identify characteristics of STIM1 cEF-hand coding variants on human disease. Most of the characterized variants are within the cEF-hand and are heterozygous autosomal dominant variants causing tubular aggregate myopathy (TAM; MIM #160565) characterized by formation of tubular membrane aggregates that cause muscle weakness (Böhm et al., 2014, 2013; Walter et al., 2015) or a more systemic Stormorken syndrome (MIM #185070) with symptoms such as TAM, miosis, and thrombocytopenia (Misceo et al., 2014; Stormorken et al., 1985). On the other hand, homozygous autosomal recessive variants can cause CRAC channelopathy and primary immunodeficiency (MIM #612783) characterized by recurrent infections, reduction in immune response, increase of lymphoproliferative disease and autoimmunity (Byun et al., 2010; Feske, 2010; Fuchs et al., 2012; Shaw and Feske, 2012).

In the cEF-hand, only one autosomal recessive variant L74P has been characterized so far, with the patient avoiding symptoms of immunodeficiency, but instead showing signs of recessive enamel development and hypohidrosis (Parry et al., 2016). We focused our analyses on missense variants possible through single nucleotide variation (SNV), as these are the variants most likely to be seen in people. We found that STIM1 missense variants possible through SNV were largely devoid of the lowest survival scores observed in our datasets (**Figure 8A**).

**Figure 8.**
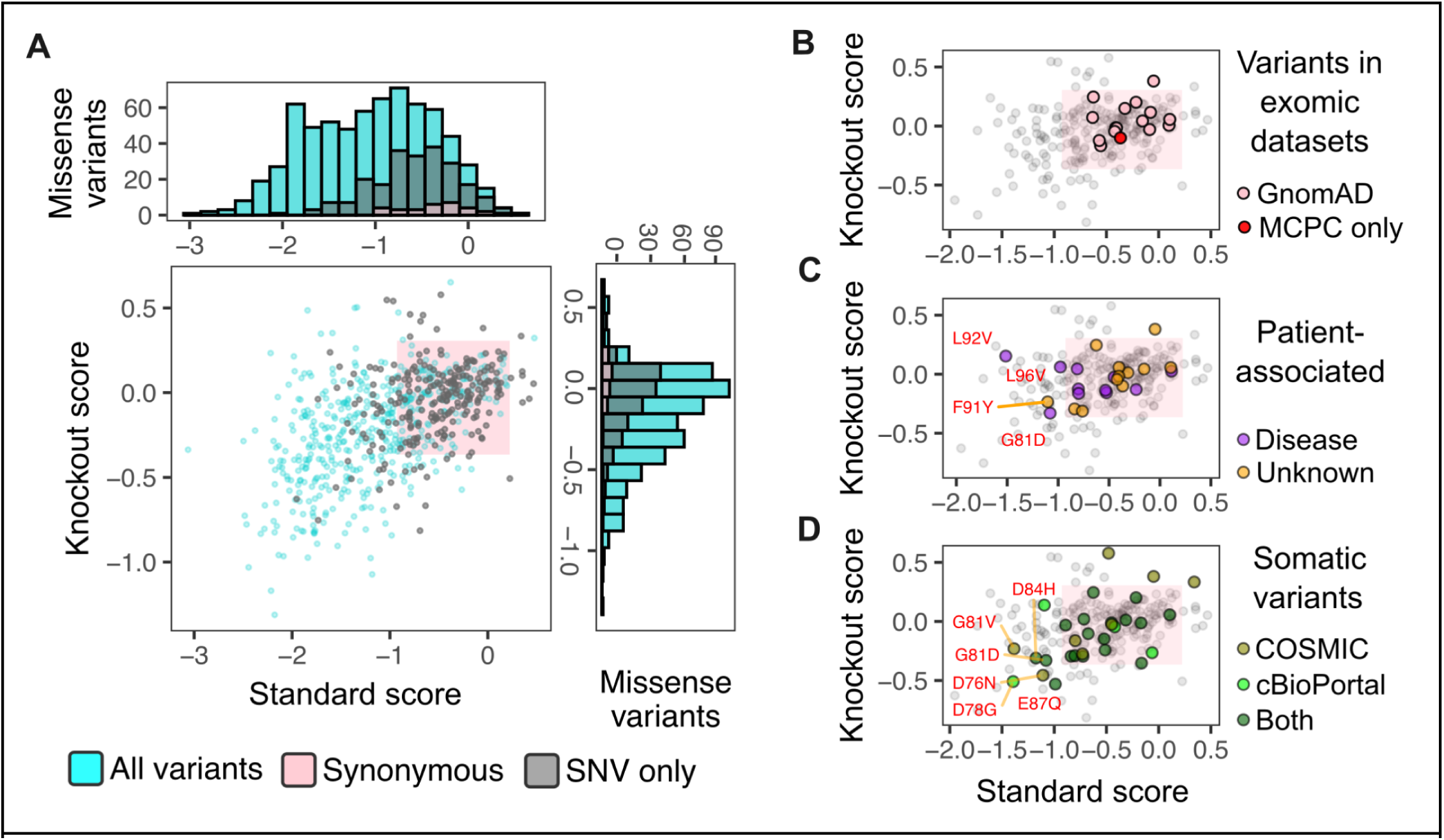
STIM1 cEF-hand variants in human disease: (A) Scatterplot and histograms of standard and knockout scores, with variants possible by single nucleotide change shown in gray. Scatterplot of mammalian survival scores for: (B) Variants observed in GnomAD or the MCPC datasets of healthy individuals. (C) Variants reported in Clinvar or in published reports describing patient clinical presentation. Magenta colored points are variants identified in patients that presented with myopathy or other STIM1-associated physiological phenotype. (D) Somatic STIM1 variants reported in COSMIC or in various cancer genomics datasets retrieved via cBioPortal. The pink boxes represent the 95 percent interval of the synonymous variant scores, defining the range of WT-like survival scores.

STIM1 cEF-hand coding variants observed in healthy humans should have few if any highly penetrant, disease-causing STIM1 variants. Indeed, no STIM1 cEF-hand missense variants observed in GnomAD exhibited scores lower than the 97.5% of the 30 synonymous variants tested in our library (**Figure 8B**). Comparatively, STIM1 cEF-hand coding variants observed in patients with myopathy have a higher likelihood of being disease-causing. We identified 21 variants within the STIM1 EF-hand listed in ClinVar, and 2 additional variants described only in published case reports. 11 of these variants are listed as pathogenic or likely pathogenic, and only 8 were fully within the synonymous range. Myopathy variants G81D, L92V, and L96V all had scores below the synonymous range in standard conditions, whereas they exhibited scores within the synonymous range in knockout conditions (**Figure 8C**).

Of the 12 variants of uncertain significance listed in ClinVar, only F91Y exhibited a score outside of the synonymous range (**Figure 8C**). Thus, while our assay results are insufficient for use in clinical interpretation, the low score exhibited by F91Y suggests it may be disease causing. Similarly, there are 47 variants of the STIM1 cEF-hand that exhibit scores lower than the synonymous distribution but have not yet been observed in sequenced patients. Our dataset suggests that these variants are likely to be disruptive and likely causative for STIM1 associated disease.

Somatic STIM1 variants are also sporadically observed in cancers, with a subset of observed variants likely contributing to cancer development. There were 30 somatic STIM1 cEF-hand variants currently observed in tumor sequencing databases, of which 7 exhibited scores below the synonymous distribution (**Figure 8D**). This included D76N, D78G, and E87Q, which had scores below the synonymous range in both cell backgrounds, whereas S64R, G81V, G81D, and D84H only had scores below the synonymous range in standard conditions. Altogether, low-scoring STIM1 variants are rarely observed, but are associated with disease whenever they are.

### Grouping STIM1 disease-associated variants by cell effect

Our finding that most disease-associated variants exhibit WT-like survival scores suggest that other assays will be needed to distinguish disease causing variants from benign ones. Furthermore, our work suggested that different STIM1 variants, both within and outside of the cEF-hand, are likely to have distinct outcomes on cell phenotype, but it is unclear which cellular mechanisms are disrupted by each variant. We thus tested a small panel of diverse STIM1 pathogenic variants, and compared the survival scores to more targeted measurements of cellular function.

We first assessed dysregulated calcium influx during SOCE (Silva-Rojas et al., 2020). We expressed WT STIM1 and four variants along with the cytoplasmic fluorescent biosensor CaMPARI2. CaMPARI2 photo-converts from green to red fluorescence emission in the presence of high calcium and 405 nm light (Moeyaert et al., 2018). Addition of the GPCR agonist carbachol resulted in high red:green ratio for TAM-associated D84G, and STRMK-associated R304W. We observed a strong inverse correlation to the standard LP scores (**Figure 9A**), consistent with cells with abnormally high calcium influx experiencing decreased survival during culture.

**Figure 9.**
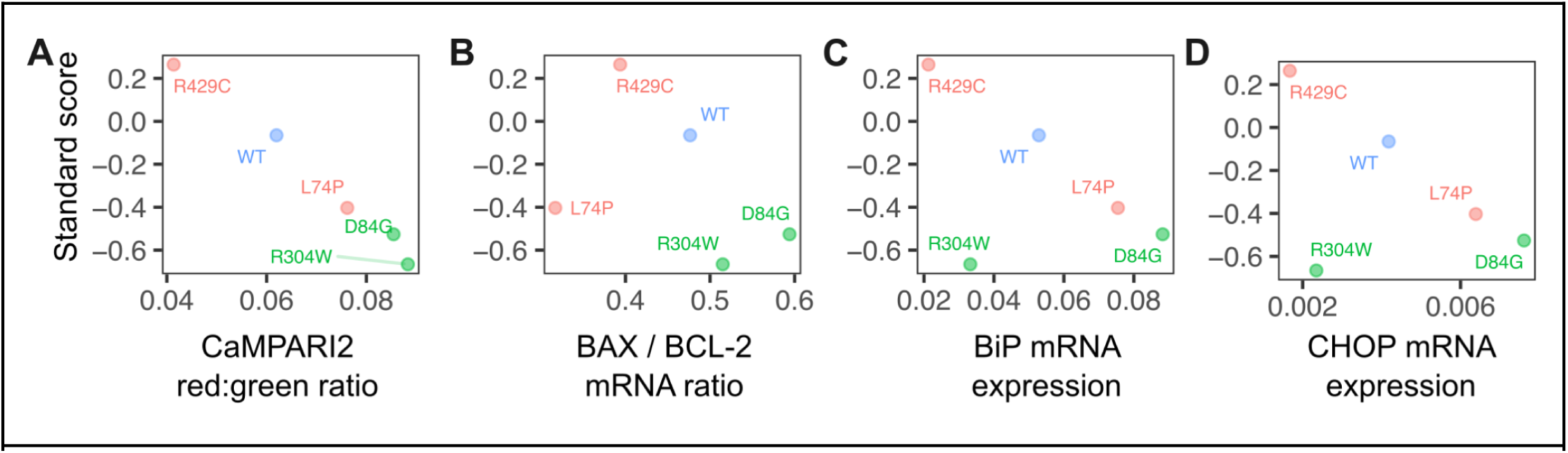
STIM variant-specific induction of cell stress pathways: (A) Scatterplot of standard LP cell survival score and calcium influx measured by CaMPARI2. (B) Scatterplot of standard LP cell survival score and mRNA expression ratio of BAX / BCL-2. (C) Scatterplot of standard LP cell survival score and BiP mRNA expression. (D) Scatterplot of standard LP cell survival score and CHOP mRNA expression. Variants in red cause immunodeficiency and variants in green cause TAM/STRMK.

Excessive calcium influx may cause calcium overload and induce various cell death pathways, including apoptosis (Danese et al., 2021). We assessed transcriptional priming for apoptosis via mitochondrial outer membrane permeabilization by quantitating the ratio of BAX and BCL-2 transcripts (**Figure 9B**). BAX/BCL-2 ratios inversely correlated with survival score, albeit with much higher variance than the CaMPARI2 measurements.

STIM1 variants that dysregulate the ER luminal environment with mistrafficking, misfolding, or unresolvable aggregation may activate ER stress pathways and contribute to disease (Wang and Kaufman, 2016), either indirectly through potential regulatory interactions with ER stress components like IRE1 (Carreras-Sureda et al., 2023), or directly as a misfolded client for an ER chaperone like BiP/HSPA5. Accordingly, muscle tissue isolated from STIM1 D84G heterozygous mice exhibited elevated ER stress transcripts relative to their wild-type littermates (Bryson et al., 2024). We thus assessed whether ER-stress pathways were induced in our variant-expressing cells.

ER-stress induces expression of BiP (Kozutsumi et al., 1988), and if unresolved, induces CHOP expression and subsequent apoptosis (Oyadomari and Mori, 2004). BiP and CHOP transcript levels both inversely correlated with survival score, with the exception of R304W, which is a cytoplasmic residue substitution less likely to impact the ER lumen (**Figure 9C, D**). L74P exhibited slightly elevated BiP and CHOP expression, similar to its CaMPARI2 fluorescence level. Notably, while generally considered loss-of-function, its mode of inactivation and associated clinical presentations (Parry et al., 2016; Rice et al., 2019) may be distinct to that of R429C (Fuchs et al., 2012; Maus et al., 2015). Taken together, different STIM1 variants may distinctly perturb downstream cellular pathways, and results from orthogonal assays can likely be leveraged to functionally distinguish STIM1 variants from one another to better link molecular effects with disease presentation.

## Discussion

We performed a deep mutational scan of the ER calcium-sensing STIM1 canonical EF-hand domain for variant overexpression induced cytotoxicity in various cell environments, yielding survival scores for 706 STIM1 coding variants. This dataset enabled functional comparisons between the three segments of secondary structure present in the cEF-hand, revealing different patterns of mutational tolerance unique to each segment. Our results suggested distinct but interrelated roles for each part of the STIM1 cEF-hand domain, particularly for the structurally unresolved calcium-free state of STIM1. Finally, we used our survival scores to characterize variants recorded in patient genome sequence databases and began to determine perturbations to cellular mechanisms caused by different disease-causing STIM1 variants. Our results provide a unique look into the molecular genetics of a specialized derivation of the commonly used calcium-binding EF-hand domain, located within an important but historically difficult to study protein, STIM1.

No single functional assay can capture all necessary information and will exhibit strengths and weaknesses worth acknowledging. The overexpression-based cell survival assay we employed captured the effects of STIM1 dysfunction on cell survival, a holistic effect not capturable with targeted assays. Unfortunately, this holisticity impedes inference of the mechanistic root causes of the observed effects. The mitochondria buffers calcium ions that enter the cytosol as a result of SOCE and uncontrolled calcium influx can result in mitochondrial calcium overload (Tanwar et al., 2021) and could be the underlying mechanism of cellular dysfunction for a subset of STIM1 variants, including R304W. Alternatively, prolonged unresolved ER stress is a possible mechanism of cell death, as misfolded proteins within the ER result in the activation of ER stress pathways and the unfolded protein response (Wang and Kaufman, 2016), which can trigger apoptosis. This mechanism is likely particularly relevant with STIM1 cEF-hand domain variants, such as D84G.

Furthermore, we cannot determine whether each low survival score variant actively triggered cell death or if it simply disturbed homeostatic cell metabolism resulting in slower growth kinetics compared to cells with WT STIM1. A panel of assays – some holistic for cell death and others more targeted such as measurement of calcium influx and ER stress gene expression – will be needed to accurately understand the mutational genotype-phenotype map for almost any protein, and especially one with the molecular complexity of STIM1 and its conformational regulation. In this paper we begin to determine the cause of cell death for a handful of known STIM1 pathogenic variants (**Figure 9A-D**).

While our recombinase-based transgenic overexpression system was necessary to perform the large-scale genetic screen, there are certain biological limitations with its use as a model system. Despite our attempts to limit the amount of STIM1 protein being expressed from our transgenic locus, the dosage of resulting STIM1 remained in vast excess of what is normally present from the endogenous copies (**Figure 2B - figure supplement 1B-C**). This is particularly relevant for STIM1, as its intracellular concentration likely alters the magnitude and kinetics of oligomerization that occurs. For many variants, this may exaggerate the phenotypic effects that would happen if only the endogenously encoded copies were altered. Regardless, the approach and its ensuing dataset yielded information useful for understanding this enigmatic protein.

Low scoring variants are likely to be detrimental to cell function across contexts. For example, retransformation of the plasmid DNA library revealed a positional depletion of variants in *E.coli* that corresponded closely with positions in the cEF-hand critical for coordinating calcium (Asp76 and Asp78) or in the central FLRE residues in helix α2 likely critical in keeping STIM1 unaggregated. Any spurious STIM1 polypeptide fragments made in bacteria will not exhibit natural localization and function. While the quantitative impacts of detrimental ectopic expression of STIM1 peptide fragments are difficult to deconvolute from the plasmid retransformation dataset, the concordance of established structurally critical residues to positions of greatest depletion suggest a common element behind both phenomena.

Low-scoring STIM1 variants were absent in healthy mammalian genomes. There are roughly three times fewer predicted loss-of-function variants observed in generally healthy human exomes aggregated in GnomAD than predicted based on chance, implying purifying selection of this gene in humans. Accordingly, the STIM1 cEF-hand is evolutionarily conserved, with all mammals exhibiting only two or fewer amino acid differences from the human sequence in the 36-residue cEF-hand. This included single amino acid differences E66D, R69C and N70S, which all exhibited WT-like scores in our synonymous distribution range (**Figure 7E**). Puzzlingly, we observed a biased lack of low score variants in our subset of STIM1 variants possible through single nucleotide change (**Figure 8A**); it is unclear whether this is by chance or selected for through an unidentified evolutionary mechanism.

Low scoring variants corresponded to those causing disease in humans. None of the 15 unique human STIM1 cEF-hand variants observed in healthy human exome databases exhibited scores outside of the synonymous variant range (**Figure 8B**). In contrast, three of the eleven known pathogenic or likely pathogenic variants exhibited low scores (**Figure 8C**). Despite its low sensitivity, the high specificity of low survival score variants associating with disease suggests that our results can be helpful for interpreting STIM1 variants in roughly ten percent of cases. We observed even more low-scoring STIM1 somatic variants in sequenced tumors. Roughly 23 percent of unique STIM1 variants exhibited scores below the synonymous variant range (**Figure 8D**). STIM1 overexpression is believed to contribute to progression of various cancers (Chen et al., 2011). Our results support a subset of tumors achieving similar outcomes though mutation of key cEF-hand residues.

There are remaining questions that require additional experiments to answer. Nonsense variants are oftentimes considered controls for DMS data analysis, as they are expected to be inert in function and the resulting phenotype would be similar to WT. In all three of our datasets, we observed a spread of scores for the nonsense variants that seemed to be largely position dependent. There is an absence of STIM1 nonsense variants in human exomes, and none observed yet in the STIM1 cEF-hand. The segment of STIM1 between the signal peptide and cEF-hand remains largely enigmatic, although multiple independent reports suggest that it plays a role in regulating STIM1 activation kinetics (Hawkins et al., 2010; Sirko et al., 2022; Zhou et al., 2009; Zhu et al., 2018). Thus, the various polypeptide fragments created by different STIM1 cEF-hand nonsense variants may each alter ER proteasis and STIM1 regulation to different extents. Further work will be needed to determine if any of these observations are related.

Large-scale mutational scans in mammalian cells is a burgeoning genetic technique, coming with its own unique pitfalls. For example, while our plasmid construct was purposefully devoid of a mammalian promoter, it inadvertently encoded a cryptic bacterial promoter (**figure supplement 4A**). These unexpected results in *E.coli* highlight the risk of assuming that variant libraries designed for mammalian expression will exhibit little to no selection upon propagation in bacteria. Similar phenomena have affected expression of some reference human cDNAs such as SCN1A (DeKeyser et al., 2021). Our results suggest that variant libraries of some proteins may be affected in a positional or variant-specific manner, which was particularly observable in our library since each position was originally prepped and quantitated separately, thus enabling roughly even variant starting frequencies that changed upon library retransformation.

Despite the challenges, large-scale genetic studies on STIM1 and similar proteins remain important. Non-globular, membrane-integral, dynamically aggregating, and potentially cytotoxic proteins like STIM1 are difficult to study with most traditional laboratory methods. Accordingly, despite intense effort, major aspects to their function elude detailed understanding. Computational approaches are rarely developed or trained for these proteins due to their relative absences from large-scale training data. Genetic and genomic approaches can be informative, but will require careful synthesis with existing information to help identify and resolve our gaps in understanding for these proteins.

## Materials and Methods

### Standard recombinant DNA generation

Recombinant DNA constructs were created with Gibson Assembly. DNA fragments were generated by amplifying 40 ng of plasmid DNA with 0.33 μM each of forward and reverse primers and 2x Kapa HIFI HotStart ReadyMix (Roche). The mixture was initially denatured at 95°C for 5 mins, and then cycled nine times at 98°C for 20 secs, 65°C for 15 secs and 72°C for 8 min, followed by a final 72°C incubation for 5 mins. Template DNA was digested through addition of 1 μL, corresponding to 20 U, of DpnI enzyme (New England Biolabs), incubated for 2 hrs at 37°C. The reactions were cleaned using Zymo clean and concentrator kit (Zymo Research) and resulting 2 μL of eluted product was mixed with 2 μL of GeneArt Gibson Assembly enzyme (Thermo Fisher, A46629), and incubated at 50°C for 30 to 60 mins. The resulting mixture was transformed into *E.coli* strain DH10β (New England Biolabs) that were rendered chemically competent with 100 mM calcium chloride, and plated overnight on LB-agar plates with 150 μg/mL ampicillin. Bacterial colonies were screened by initial overnight growth in LB media supplemented with 150 μg/mL ampicillin, followed by plasmid extraction using a GeneJet miniprep plasmid kit (Thermo Fisher). For each candidate clone, successful insertion of transgene into the template plasmid was verified through Sanger sequencing (Applied Biosystems 3730 genetic analyzer). Whole plasmid sequencing was performed for all selected clonal variants used in the study (Plasmidsaurus, Oxford Nanopore Technologies). For **Figure 2D**, variant plasmids were PCR amplified again with forward and reverse primers containing known 10 nucleotide sequences, together forming a double barcode unique to each variant (**Supplement Table 2**).

The following plasmids from Addgene were used as template material in the plasmid constructs generated in this work: pLNCX2-STIM1 was a gift from Shengyu Yang (Addgene plasmid # 89817; http://n2t.net/addgene:89817; RRID:Addgene_89817) (Sun et al., 2014); pcDNA3.1-mGreenLantern was a gift from Gregory Petsko (Addgene plasmid # 161912; http://n2t.net/addgene:161912; RRID:Addgene_161912) (Campbell et al., 2020); pAAV_hsyn_NES-his-CaMPARI2-F391W-WPRE-SV40 was a gift from Eric Schreiter (Addgene plasmid # 101061; http://n2t.net/addgene:101061; RRID:Addgene_101061) (Moeyaert et al., 2018). The following plasmids that we generated were deposited into Addgene for future distribution: pNK036C_AttB_mkozak_mGreenLantern-STIM1_IRES_mCherry-H2A-P2A-PuroR (Addgene plasmid # 232938); J364B_AttB_mkozak_STIM1_IRES_CaMPARI2-P2A-PuroR (Addgene plasmid # 232939).

### STIM1 cEF-hand domain variant library generation

Plasmid pNK005C_BattB_mkozak_STIM1_IRES_mCherry-H2A-P2A-PuroR (Addgene plasmid # 200639) was used as the template construct for cEF-hand variant library generation. Degenerate NNK primers were constructed such that the forward primers had 18 nucleotides before the NNK on the 5’ end and 15 nucleotides after on the 3’ end. The reverse primers were 25 nucleotides before the NNK. They were created using SSM_primer_EFhand.py (https://github.com/MatreyekLab/STIM1_cEFhand) and ordered from Integrated DNA Technologies. Each position was amplified separately using the protocol detailed in ‘Recombinant DNA generation. The DNA concentration of each mutated position was measured using a NanoDrop spectrophotometer (Thermo Fisher Scientific). The final library was constructed by mixing equimolar amounts of each variant position. The final mixture was sequenced through Illumina sequencing to determine the relative frequencies of each variant in the plasmid library (**figure supplement 1C-E**).

### Cell culture and STIM1-/- cell line generation

HEK293T-derived landing pad cells were cultured in D10 media, Dulbecco’s Modified Eagle Medium (DMEM) (Corning) supplemented with 10% fetal bovine serum (Gibco), 100 U/mL Penicillin, 0.1 mg/mL Streptomycin (Corning), either alone (D10) or supplemented with 2 μg/mL Doxycycline (D10X). Cells were detached using Trypsin-EDTA (0.25%, Corning).

Two related but distinct clones of HEK 293T cells were used in this work. HEK293T single landing pad cells, also identified to as LLP-Int-iCasp9-Blast clone 3 cells (HEK293T G542Ac3), were used for all experiments where endogenous STIM1 was present (Shukla et al., 2021) and are referred to as “standard LP cells” in this study. Prior to recombination, these cells were maintained in D10X media supplemented with 10 μg/mL Blasticidin. HEK293T G542Ac3 cells were modified with a second landing pad site to yield double landing pad cells (HEK293T G542Ac3 G783Ac11), and were maintained in D10X media supplemented with 10 μg/mL Blasticidin and 50 μg/mL Hygromycin (Roelle et al., 2023).

For this study, double landing pad cells were knocked out for endogenous full-length STIM1 using CRISPR-Cas9. Cas9 sgRNAs targeting the introns flanking STIM1 exon 2 were created by modifying plasmid gRNA_AAVS1-T2, which was a gift from George Church (Addgene plasmid # 41818; http://n2t.net/addgene:41818; RRID:Addgene_41818). The sgRNA targeting sequences corresponded to GGGAGGGAGAGAGATCCCTG in G1123C_KAN_Stim1-GGG-guide, and CTGTCACTGTACAAGTAGCC in G1124A_KAN_Stim1-GGC-guide. These plasmids were mixed with the Cas9 expression vector, PX458, and transfected using Xfect transfection reagent (Takara). pSpcas9(BB)-2A-GFP (PX458) was a gift from Feng Zhang (Addgene plasmid # 48138; http://n2t.net/addgene:48138; RRID:Addgene_48138). Two days later, the cells were visually confirmed for high green fluorescence, and replated at low density into 100 mm plates, with D10X supplemented with 100 μg/mL zeocin. Two weeks later, single colonies of cells were picked and transferred to 6-well plates to grow. Genomic DNA from isolated colonies were extracted and amplified using primers flanking exon 2 of STIM1. The reactions were fractionated by agarose gel electrophoresis and the presence of amplicon bands smaller than those present in unmodified cells indicated absence of the targeted exon. Further confirmation was performed through western blot (**Figure 1D**). These cells are referred to as “knockout LP cells” in this manuscript.

### Recombination of plasmid into landing pad cells

For recombination of clonal control plasmids, 600,000 cells were plated per well of a 6 well plate in D10X media a day before transfection. For each plasmid, 5 μg of DNA was combined with 100 μL of Xfect Reaction Buffer and 1.5 μL of Xfect Polymer (Takara) and added to cells. Prior to recombination, the landing pad encodes iCasp9, an engineered caspase, permitting inducible depletion of unmodified cells (Roelle et al., 2023; Shukla et al., 2021). 10 nM AP1903 (Apex Bio) was added 72 hours after transfection to negatively select for recombined cells. Fresh D10X media was added the next day and recombined cells are allowed to grow to ∼70% confluence before adding Puromycin to a final concentration of 1 μg/mL to generate a pure population of recombined cells for downstream applications.

Prior to performing recombinations for the library-based experiments, we performed a statistical simulation which indicated that at least 300,000 recombined cells were needed for every variant in the library to have independently recombined into 10 different cells (**figure supplement 2D**). To achieve this with a relatively conservative estimate of 1% recombination efficiency meant that we needed to transfect approximately 3 million cells. Thus, for each replicate experiment, standard or knockout LP cells were plated on three 100 mm plates with 3.5 million cells per plate a day before transfection. 30 μg of the variant library plasmid mixture was combined to a final volume of 600 μL of Xfect Reaction Buffer and 9 μL of Xfect Polymer (Takara) and added to the cells growing in D10X. STIM1 knockout was performed in double landing pad cells and to simplify library recombination in these cells, the cells were first recombined with plasmid G1095C_AttB(GA)_jGCaMP7c_IRES_shBleR-P2A-HygroR (Addgene # 200638), and selected for successful integrants of the second attP[GA] landing pad site with final concentrations of 10 μg/mL blasticidin and 100 μg/mL zeocin. This ensured that only a single site was available for recombination of the STIM1 cEF-hand library. 10 nM AP1903 (Apex Bio) was added 72 hours after transfection to negatively select for recombined cells. Fresh D10X media was added the next day. Cells were sampled 5, 7 and 9 days after transfection.

### Flow cytometry and analysis

Recombined landing pad cells were detached using Trypsin-EDTA and resuspended in D10X media. Flow cytometry was performed on the Attune NxT (Thermo Fisher Scientific). Cells expressing mCherry were excited using a 561 nm laser and emission was detected through a 620/15 nm bandpass filter. BFP was excited using a 405 nm laser and emission was detected through a 440/50 nm bandpass filter. FlowJo version 10.8.0 was used for analysis of flow cytometry data. Live cells were gated using FSC-A and SSC-A and further gated for singlets using FSC-A and FSC-H. Singlet cells were then examined for mCherry and BFP fluorescence, with the subpopulation of cells exhibiting high mCherry but low BFP fluorescence representing the recombined cell population. The proportion of the singlet gated cells that were found in this subpopulation was used to monitor the relative cell survival following clonal transfections of control STIM1 variants.

### Western Blot

Standard LP cells, knockout LP cells, standard LP cells with overexpressed WT STIM1 (pNK005C) and knockout LP cells with overexpressed WT STIM1 were harvested and lysed with 1x RIPA buffer (Thermo Fisher Scientific, 89901) containing 1:100 concentration of protease inhibitor cocktail (Thermo Fisher Scientific). Protein lysates were quantified with BCA assay (Thermo Fisher Scientific) and calculations were done to load 60 μg of protein to detect endogenous STIM1 in standard LP cells and knockout LP cells or 20 μg for overexpressed STIM1 in standard and knockout LP cells. Lysates were denatured using LDS sample buffer (Thermo Fisher Scientific) and run on a 4-12% SDS Bis-Tris gel (GenScript) with MOPS running buffer (GenScript). Proteins were transferred to 0.2 μM PVDF membrane (Thermo Fisher Scientific) and stained for total protein using Ponceau S stain (Thermo Fisher Scientific). The membrane was blocked with 5% non-fat milk in TBST and then probed for STIM1 with anti-STIM1 monoclonal antibody (CDN3H4) (1:1000 dilution, Thermo Fisher Scientific). Goat anti-mouse secondary HRP-conjugated antibody was used at 1:10000 dilution. HRP-conjugated β-actin (Thermo Fisher Scientific, 1:10000 dilution) was used as loading control. Imaging was done using an Amersham Imager 600 (GE) for **Figure 1D** and iBright FL1500 Imaging System (Thermo Fisher Scientific) for the rest of the figures.

### Genomic DNA extraction and sample preparation for NGS

Genomic DNA was extracted from cells using the DNeasy Blood and Tissue Kit (Qiagen) and concentrations were measured by Qubit Fluorometer (Thermo Fisher Scientific). PCR amplification was performed with 1.25 μM of each forward and reverse primer with 10 nucleotide long unique barcodes for sample multiplexing (**Supplement Table 2**), 100 ng of gDNA, 2x Phusion primer mix (Thermo Fisher Scientific) and nuclease-free water to a final volume of 40 μL. The PCR conditions were: 95°C for 3 minutes, repeat 95°C 15 seconds, 60°C 15 seconds and 72°C 30 seconds 35 times, 72°C for 1 minute, 4°C hold. Reactions were run on a 1% agarose gel to extract amplicon bands. DNA was extracted from the gel bands using the GeneJet Gel Extraction and DNA Cleanup Micro Kit (ThermoFisher Scientific). The DNA concentration of each sample was measured by Qubit, normalized to 5 ng/μL, and mixed in equal volumes to make the final library. Samples were run on NextSeq 550 using a Mid Output 150 cycle kit by the CWRU Genomics Core.

### DMS of select STIM1 cEF-hand positions in bacteria

The retransformed variant mini-library was made by equimolar mixing of select previously created NNK-mutagenized positions. The library was transformed into three strains of bacteria - DH10β (New England Biolabs), DH5α (Thermo Fisher Scientific) and BL21-DE3 (Thermo Fisher Scientific). For DH10β and BL21, 5 μL of plasmid library was added to the bacteria and placed on ice for 15 minutes before heatshocking at 42°C for 1 minute. For DH5α, 5 μL of plasmid library was added to the bacteria and electroporated with Gene Pulser Xcell (BioRad) in a 0.1 cm cuvette using these settings: 1.8 kv, 200 Ω, 25 μF. All conditions were grown out in 1 mL S.O.C recovery media (New England Biolabs) for 1 hour and then transferred to respective growth media for overnight incubation. The growth media used for these experiments were Luria Broth (LB, VWR), M9 Minimal Media (Thermo Fisher Scientific) supplemented with 20% D-Glucose solution and 2 mM final concentration of MgSO_4_ solution (MM), or MM with additional 100 μM final concentration of CaCl_2_ solution (Ca). Plasmid DNA was extracted from bacteria using GeneJET Plasmid Miniprep Kit (Thermo Fisher Scientific). The cEF-hand domain region of each DNA sample was amplified using forward and reverse primers with 10 unique nucleotide barcodes (**Supplement Table 2**) and samples were mixed and sequenced as stated above.

### Data analysis

Raw FASTQ files were paired using fastq-join 1.3.1 (Aronesty, 2013). A Python (3.11.4) script titled “fastq_process_improve.py” was used to parse through paired FASTQ reads for nucleotide sequence, translate into amino acid sequences, and call variants for each read. Reads with more than one variant called were filtered out. Sequencing of the three plasmid library samples was performed with a small-scale MiSeq 2×250 cycle Amp-EZ service (GeneWiz by Azenta Life Sciences). These FASTQ files were parsed using a slightly modified Python script titled “ampez_fastq.py”. For samples used in **Figure 2G**, double barcodes were used for each of the known variants and WT. Double barcodes were extracted by comparing a csv file “10ntR1_10ntR2.csv” containing double barcodes and counted using Python script “bc_process.py”.

The DMS experiments were performed a minimum of five times and the validation experiments three times. Survival ratios were calculated as ratio of the frequency of the variant in cells divided by the frequency of the variant in the plasmid library. Survival scores were calculated as the log10 transformation of the survival ratios. Survival scores for each variant in standard LP cells and knockout LP cells, and the transformation scores in bacteria, were tabulated into **Supplement Table 1**. All subsequent analysis was done in R version 4.3.1 (2023-06-16). An R markdown file with code to recreate analysis (STIM1_cEFhand_DMS.Rmd), along with the relevant data files and the python script for the initial processing of FASTQ files can be found in the MatreyekLab Github repository under “MatreyekLab/STIM1_cEFhand”. Raw FASTQ data can be found at the NCBI GEO repository under accession number GSE287849. (**Supplement Table 3**).

Germline STIM1 coding variants in generally healthy individuals were compiled after accessing the following databases: GnomAD (Chen et al., 2024) on September 24, 2024, TopMed BRAVO freeze 10 (Taliun et al., 2021), and Regeneron Mexico City Prospective Study (MCPS) (Ziyatdinov et al., 2023). Somatic STIM1 coding variants from tumor samples were compiled after accessing the following databases: cBioPortal (Cerami et al., 2012) on October 1, 2024, and COSMIC (Tate et al., 2019) on September 24, 2024. Disease-associated STIM1 germline variants were compiled from ClinVar (Landrum et al., 2014) and manual literature curation (See **Supplement Table 5**).

### Fluorescent microscopy with mGreenLantern tagged STIM1

Fluorescent microscopy was performed with plasmid pNK036C_AttB_mkozak_[mGreenLantern]-STIM1_IRES_mCherry-H2A-P2A-PuroR, and various derivatives with single amino acids mutated. In these constructs, mGreenLantern was inserted in between STIM1 residues 39 and 40 which is between the signal sequence and the cEF-hand domain, as done previously (Wu et al., 2006). The constructs were recombined into standard LP cells and were grown in D10X media for 3 days before negative selection with 10 nM AP1903. After cells reached 70% confluence, positive selection was performed with 1 μg/mL Puromycin for 7 days to generate a pure population of recombined cells. 8-well chamber slides were coated with 50 μg/mL Poly-L-lysine (Sigma) for 5 minutes and let dry for 2 hours at 37°C before cells were plated. Once cells reached 70% confluence, they were fixed with 4% formaldehyde, mounted with coverslip and ProLife Anti-fade Diamond mountant (Thermo Fisher) and left to cure for 24 hours before imaging.

Fluorescent imaging was performed on a Nikon Ti-2E fluorescent microscope, with a SOLA SM II 365 light engine (Lumencor), CFI Plan Apochromat DM Lambda 20X objective, GFP (No. 96392) and Texas Red (No. 96395) filter sets and a DS-QI2 monochrome CMOS camera. Exposure time was kept constant for all samples at 100 ms for GFP and 75 ms for Texas Red.

### Calcium influx measurements with CaMPARI2

Calcium influx measurements were performed with plasmid J364B_AttB_mkozak_STIM1_IRES_CaMPARI2-P2A-PuroR, and derivatives encoding various STIM1 variants. The constructs were recombined into standard LP cells and were grown in D10X media for 3 days before negative selection with 10 nM AP1903. After reaching 70% confluence, positive selection was done with 1 μg/mL Puromycin for 7 days to generate a pure population of recombined cells.

CaMPARI2 expressing cells were trypsinized and plated on black-walled, clear and flat bottomed 96-well plates (Greiner) in D10X media containing 1.8 mM calcium with two technical replicates for untreated and treated conditions. After 24 hours, treatment conditions were pulsed with 100 μM carbachol (Fisher) in the presence of 405 nm light for 3 minutes. Cells were placed in the incubator at 37°C for 3 hours before they were measured, as this reduced background fluorescence. The fluorescence intensity of the cells were read on a BioTek SynergyH1 plate reader at 488 nm excitation and 520 nm emission for green and 560 nm excitation and 600 nm emission for red. The ratio of fluorescence intensity was calculated as mean fluorescence intensity of red:green. This experiment was repeated three separate times.

### Quantitative PCR (qPCR) for ER stress gene expression

Standard LP cells were grown out in D10 media without Dox for 2 days before beginning the experiment to reduce landing pad Bxb1 expression. Cells were recombined with STIM1 variant plasmids and 10% total DNA of exogenous Bxb1 using pCAG-NLS-HA-coBxb1 as mentioned above. pCAG-NLS-HA-Bxb1 was a gift from Pawel Pelczar (Addgene plasmid # 51271; http://n2t.net/addgene:51271; RRID:Addgene_51271) (Hermann et al., 2014). After 3 days, cells were treated with 2 μg/mL of Doxycycline and 10 ng/mL of AP1903 for negative selection. Cells were harvested 24 hours later. mRNA was extracted from the cells using an RNeasy mini kit (Qiagen). 1 μg of mRNA was converted into cDNA using High-Capacity cDNA Reverse Transcription Kit (Thermo Fisher) and 1 μL of the resulting cDNA was used to set up qPCR reactions using SYBR Green PCR Master Mix (Thermo Fisher) with primers for BCL-2, BAX, BiP and CHOP (**Supplement Table 4**). qPCR reactions were performed on a BioRad CFX Opus 96 Real-Time PCR System. Actin was used as the housekeeping gene and all Cq values were normalized to actin Cq values. The equation used to calculate gene expression was 2^-ΔCq^ where ΔCq denotes the Cq of the gene of interest subtracted from the Cq of actin) (Schmittgen and Livak, 2008).

### Supporting information

Supplementary Figures 1-6 are included in a separate PDF file

Supplementary Tables 1-5 are included in a xslx files.

## Acknowledgements

The authors wish to thank Simone Edelheit and Milena Zelembaba of the Genomics Core Facility of the CWRU School of Medicine’s Genetics and Genome Sciences Department. The Cytometry & Imaging Microscopy Shared Resource of the Case Comprehensive Cancer Center was supported by NIH grants P30CA043703 and S10OD021559.

## Additional Information

### Funding

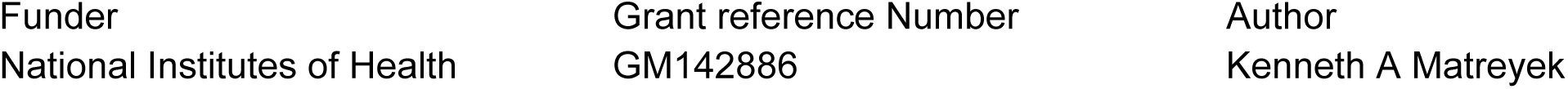

The funders had no role in study design, data collection and interpretation, or the decision to submit the work for publication.

### Author contributions

Nisha D Kamath, Conceptualization, Methodology, Investigation, Validation, Data curation, Formal analysis, Visualization, Writing - original draft, Writing - review and editing; Kenneth A Matreyek, Conceptualization, Methodology, Investigation, Data curation, Formal analysis, Visualization, Writing - review and editing, Supervision, Funding acquisition, Methodology, Project administration.

