## Supplement Figures for "Comprehensive mutational characterization of the calcium-sensing STIM1 EF-hand reveals residues essential for structure and function"

### Additional files

### Supplement Tables:

1. Supplement Table 1: Survival scores for cell survival DMS in standard LP, knockout LP, bacteria strains, and validation experiments
2. Supplement Table 2: Primers used for NGS experiments
3. Supplement Table 3: Names of FASTQ files
4. Supplement Table 4: qPCR primers used for gene expression experiments
5. Supplement Table 5: Compilation of results from all databases accessed for this study

### Supplement Figures:

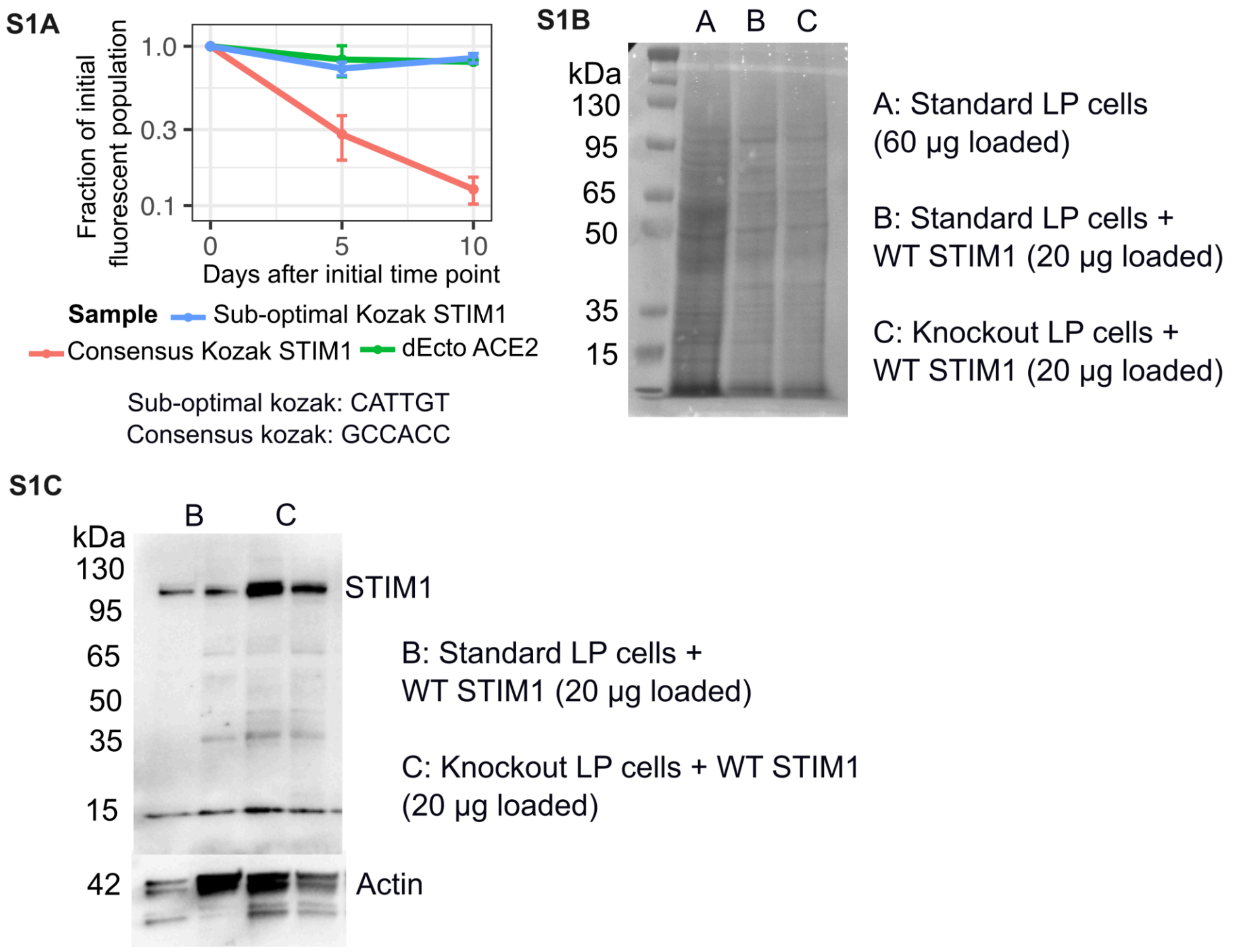


**Supplement figure 1**: A) Relative fraction of STIM1 overexpressing cells remaining in culture over time, as measured by flow cytometry for mCherry fluorescence, for STIM1 overexpression under an sub-optimal or consensus Kozak sequence. B) Total protein analysis for western blot of standard LP cells, standard LP cells with WT STIM1 overexpressed, and knockout LP cells with WT STIM1 overexpressed. C) Western blot of two replicates of overexpressed STIM1 in standard LP cells and knockout LP cells.

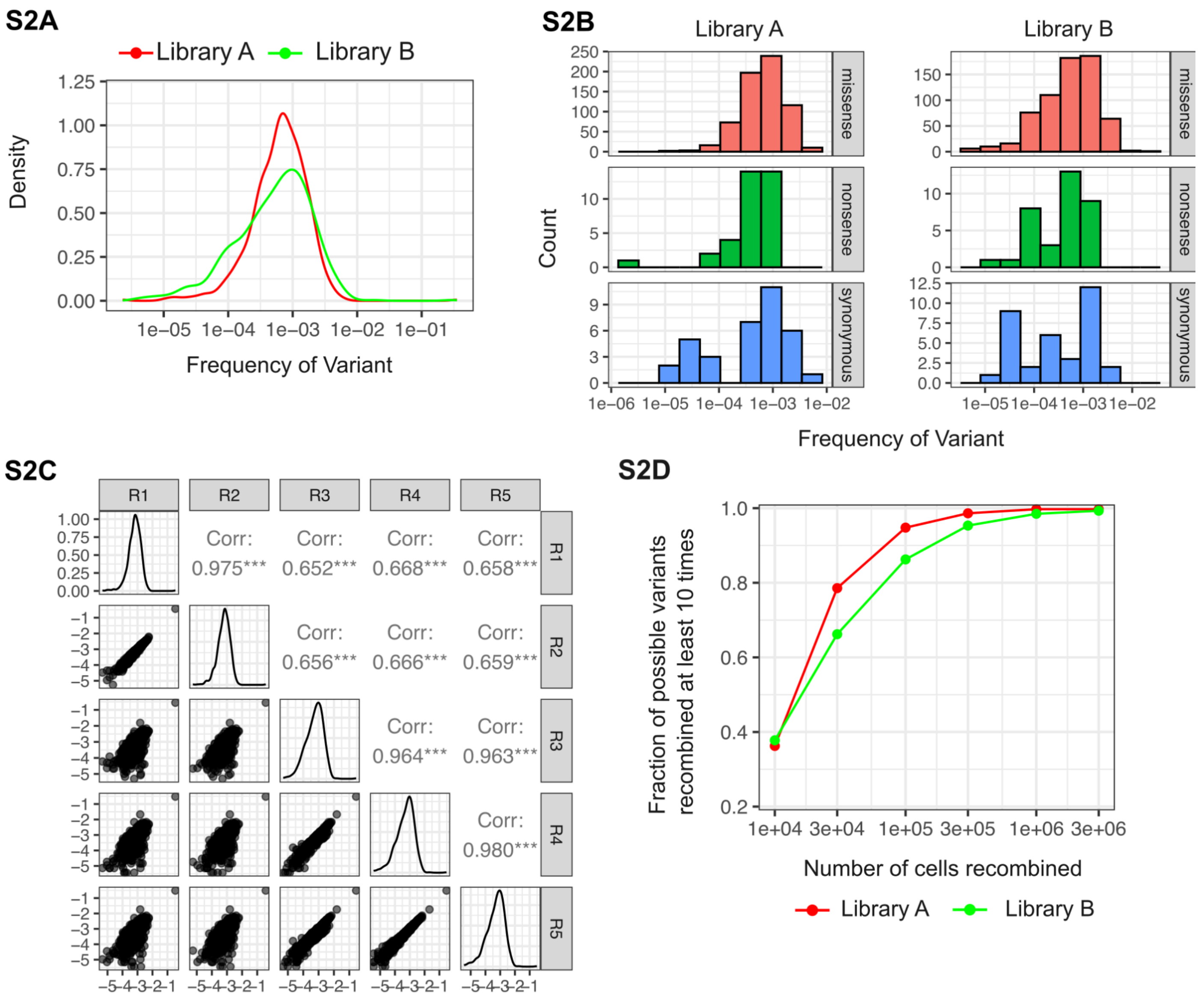


**Figure supplement 2**: A) Density plot of frequency of variants in plasmid libraries used for standard (library A) and knockout LP (library B) DMS experiments. B) Histogram of missense, nonsense and synonymous STIM1 cEF-hand variants present in both plasmid libraries. C) Correlation plot of each sequenced replicate of the plasmid library. R1 and R2 were library A, R3-R5 were library B. D) Simulation to determine the number of recombined cells needed to observe a variant at least 10 times.

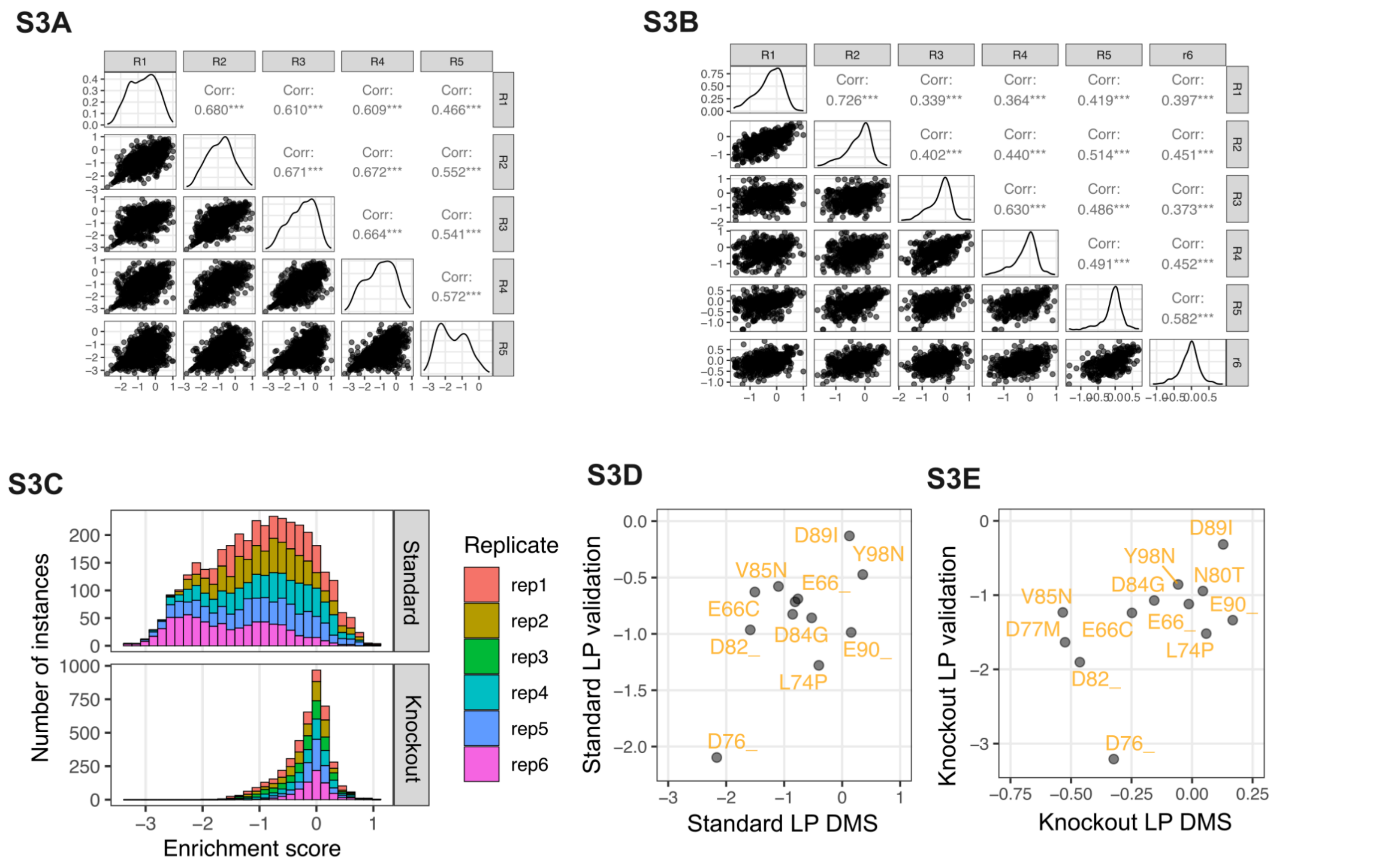


**Figure supplement 3**: A) Correlation plot of standard LP DMS replicates. B) Correlation plot of knockout LP DMS replicates. C) Histogram of variant score distributions seen in standard and knockout LP DMS, colored by replicate experiment. D) Scatterplot of DMS score vs small scale validation scores in standard LP cells. E) Scatterplot of DMS score vs small scale validation scores in knockout LP cells.

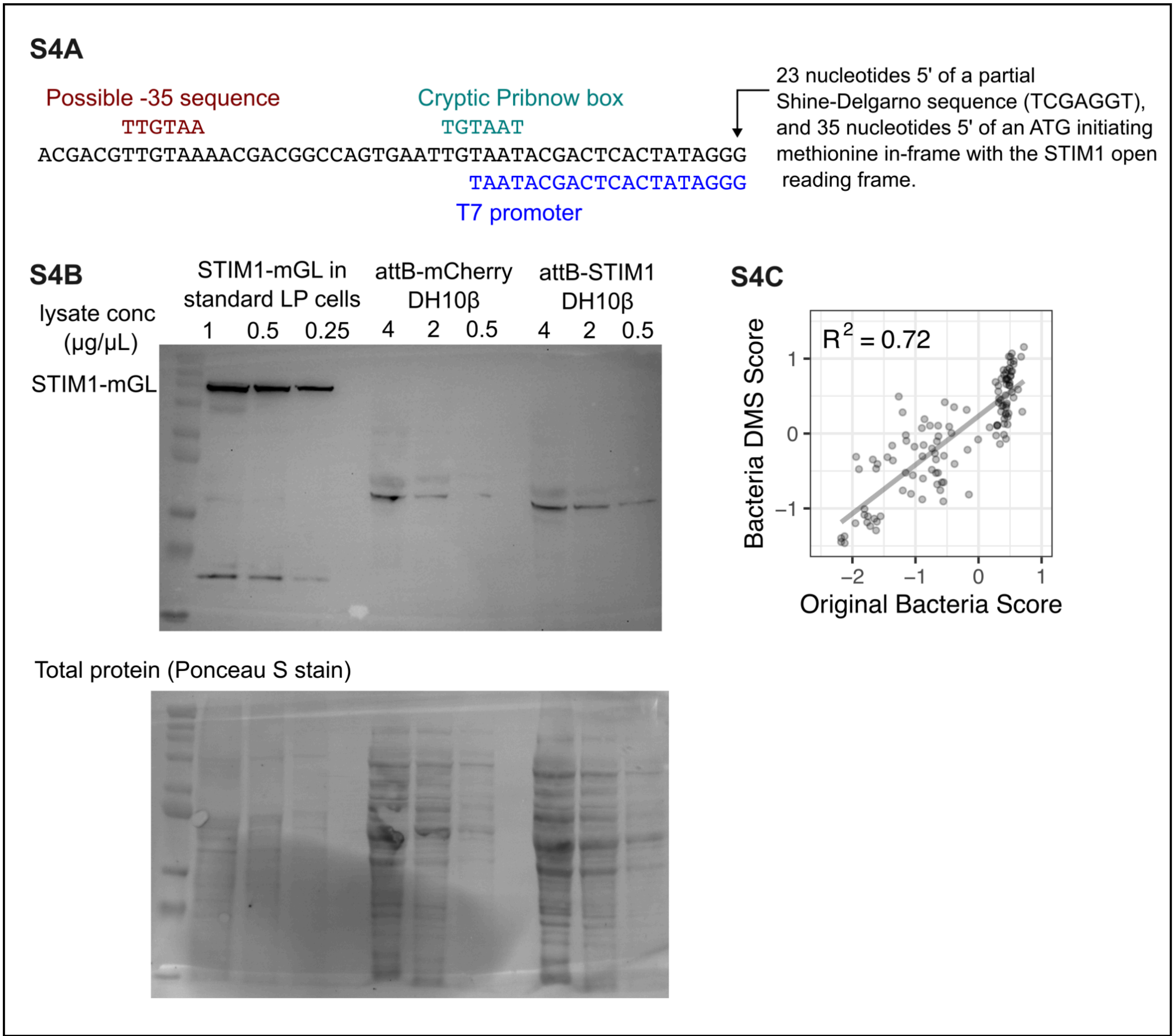


**Figure supplement 4**: A) Schematic representation of the cryptic bacterial promoter retrospectively identified in the plasmid. B) Western blot of STIM1 overexpression in DH10β bacteria. C) Scatterplot of transformation scores from first DMS to the scores from the second DMS in DH10β cells.

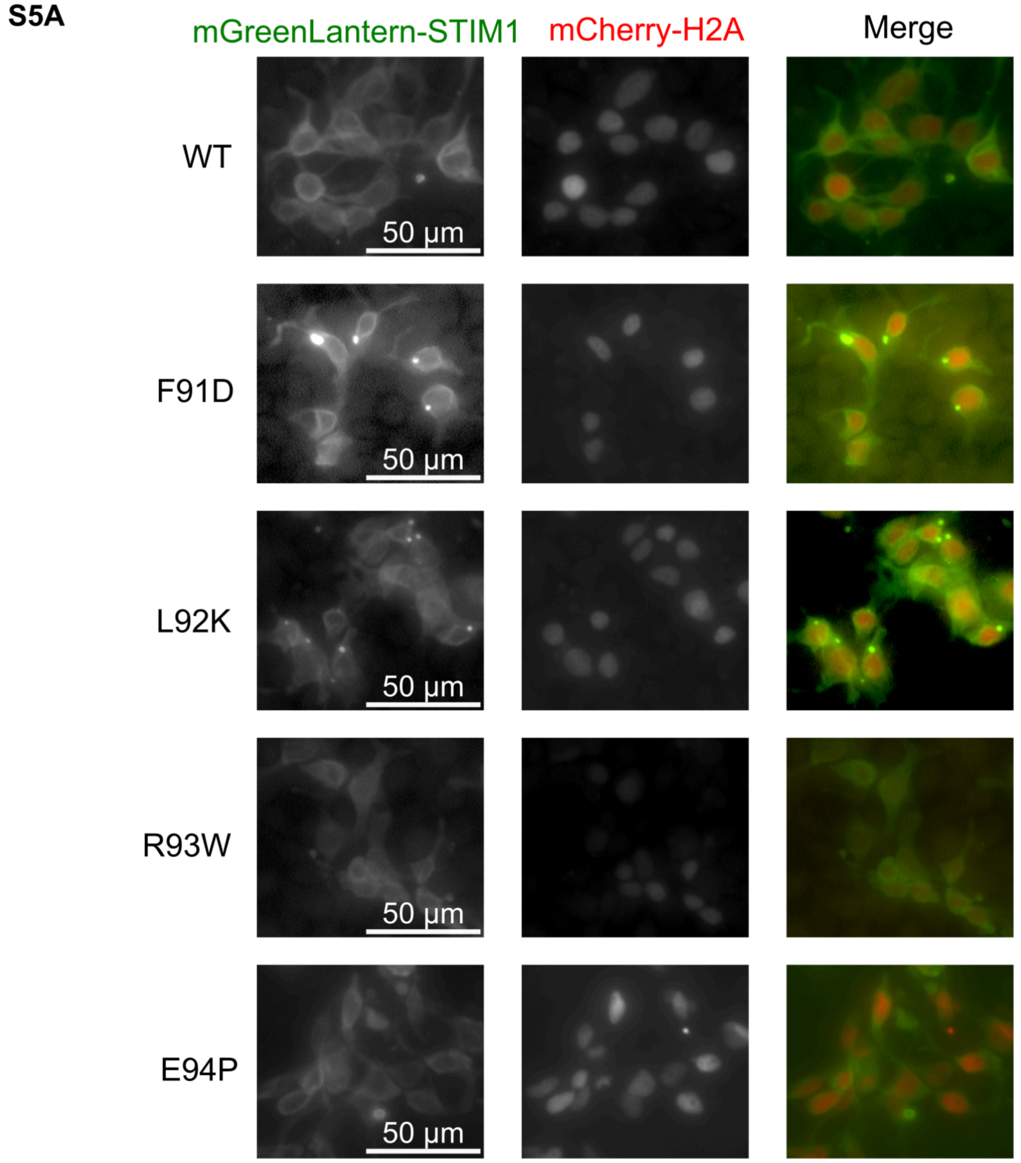


**Figure supplement 5:** A) Raw fluorescence microscopy images of mGreenLantern-tagged STIM1 variants in standard LP cells.

**S6A**

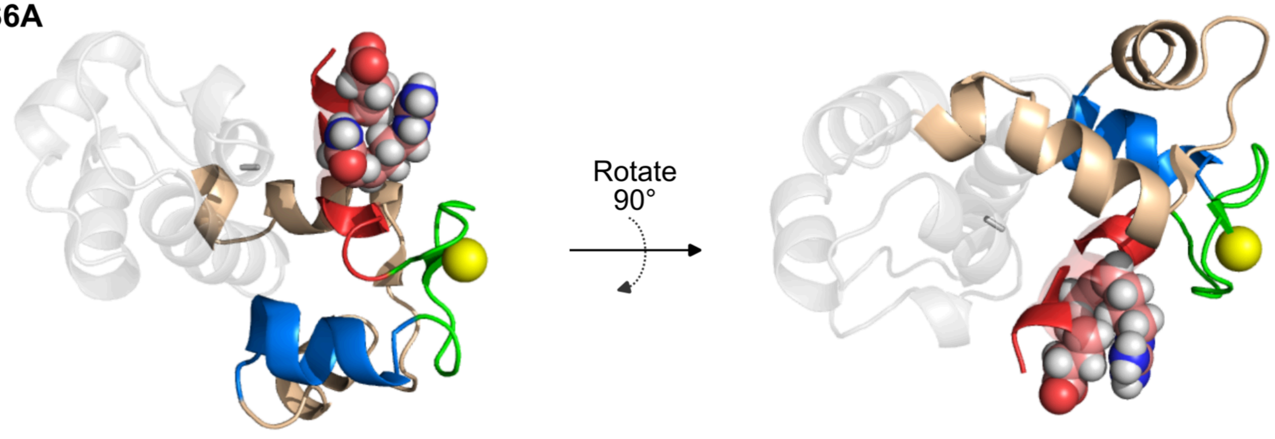


**Figure supplement 6:** Pymol visualization of the STIM1 EF-SAM calcium loaded NMR structure (PDB: 2k60), with the STIM1 cEF-hand positions differing in mammalian orthologs (Glu66, Arg69, Asn70) shown as spheres. The calcium ion is shown as a yellow sphere. Helix α1 is color red, the EF-motif is colored green, and helix α2 is colored blue. The hidden EF-hand is colored beige, whereas the SAM domain is shown in semi-transparent gray.
